# FoxO transcription factors actuate the formative pluripotency specific gene expression programme

**DOI:** 10.1101/2024.01.13.575494

**Authors:** Laura Santini, Saskia Kowald, Giovanni Sestini, Nicolas Rivron, Martin Leeb

**Affiliations:** Max Perutz Laboratories Vienna, University of Vienna, Vienna BioCenter, 1030 Vienna, Austria; Vienna BioCenter PhD Program, Doctoral School of the University of Vienna, Medical University of Vienna, 1030 Vienna, Austria; Institute of Molecular Biotechnology of the Austrian Academy of Sciences (IMBA), Vienna BioCenter, 1030 Vienna, Austria

## Abstract

Naïve pluripotency is sustained by a self-reinforcing gene regulatory network (GRN) comprising core and naïve pluripotency-specific transcription factors (TFs). Upon exiting naïve pluripotency, ES cells transition through a formative post-implantation-like pluripotent state, where they acquire competence for lineage-choice. However, the mechanisms underlying disengagement from the naïve GRN and initiation of the formative GRN are unclear. Here, we demonstrate that phosphorylated AKT acts as a gatekeeper that prevents nuclear localization of FoxO TFs in naïve ESCs. PTEN-mediated reduction of AKT activity upon exit from naïve pluripotency allows nuclear entry of FoxO TFs, enforcing a cell fate transition by binding and activating formative pluripotency-specific enhancers. Indeed, FoxO TFs are necessary and sufficient for transition from the naïve to the formative pluripotent state. Our work uncovers a pivotal role for FoxO TFs and AKT signalling in mechanisms establishing formative post-implantation pluripotency, a critical early embryonic cell fate transition.

## INTRODUCTION

Pluripotent cells can give rise to all specialised cells that form an adult organism. During mouse embryonic development, a population of naïve pluripotent cells arises in the pre-implantation epiblast around embryonic day E3.5-E4.5 (Boroviak et al. 2014). During the transition from pre- to post-implantation development, epiblast cells navigate through a continuum of pluripotency states. Starting from a naïve state with unrestricted potential, cells transit through a formative state to acquire competence for somatic and germ cell lineage specification, ultimately entering a primed state where they initiate expression of lineage markers (Kinoshita et al. 2021; Kinoshita and Smith 2018; Nichols and Smith 2009; A. Smith 2017).

The naïve pluripotent state of pre-implantation epiblast cells can be captured *in vitro* using mouse embryonic stem cells (mESCs) (Evans and Kaufman 1981; Martin 1981). Maintenance of a homogeneous ground-state of pluripotency requires the addition of two small molecule inhibitors to the culture media: PD0325901 (MEK1/2 inhibitor) and CHIRON (GSK3ɑ/ꞵ inhibitor), collectively referred to as 2i (Ying et al. 2008). mESCs cultured in 2i resemble the E4.5 pre-implantation epiblast in terms of epigenetic and transcriptional status (Boroviak et al. 2014, 2015; Ficz et al. 2013; Lee, Hore, and Reik 2014). Naïve identity is defined by the expression of a self-reinforcing gene regulatory network (GRN) that comprises the core pluripotency transcription factors (TFs) *Oct4* (gene name: *Pouf51*) and *Sox2*, and naïve-specific TFs including *Nanog*, *Esrrb*, *Klf4,* and others (Chen et al. 2008; Dunn et al. 2014; Martello and Smith 2014; Niwa et al. 2009).

A balanced interplay of several signalling inputs is responsible for the maintenance of the naïve-specific GRN (Huang et al. 2015). The cytokine LIF (Leukaemia Inhibitory Factor) plays a key role in the GRN and was the first identified exogenous factor that can support mouse ESC culture along with serum supplementation (Smith et al. 1988; Williams et al. 1988). LIF mainly activates the JAK/STAT3 and PI3K/AKT pathways which are crucial to sustain naïve pluripotency (Niwa et al. 2009). Although the role of the JAK/STAT pathway has been intensively studied, the function of PI3K/AKT signalling in pluripotency has received much less attention (Ohtsuka, Nakai-Futatsugi, and Niwa 2015). Overexpression of a constitutively active form of AKT is sufficient to maintain mESCs in an undifferentiated state, even in the absence of LIF (Watanabe et al. 2006). PI3K/AKT signalling is thought to support naïve pluripotency through inhibition of both the MEK/ERK and the GSK3 pathways (Paling et al. 2004; Wang et al. 2020; Wray et al. 2011), although the underlying mechanisms are unclear. Furthermore, AKT signalling feeds directly into the naïve GRN by activating the expression of *Tbx3* and *Nanog* (Niwa et al. 2009; Storm et al. 2007).

Exit from the naïve pluripotent state and initiation of formative pluripotency can be recapitulated *in vitro* by releasing cells from 2i inhibition into basal N2B27 medium. This change in conditions leads to loss of self-renewal in the naïve state and an irreversible commitment to differentiate approximately 48h after 2i withdrawal (Wray et al. 2011). The exit from naïve pluripotency results in the dismantling of the naïve-specific GRN and the establishment of a new formative state-specific GRN. This is accompanied by a profound shift in the signalling landscape: LIF and AKT signalling are reduced, concomitant with an increase in FGF/ERK activity (Kunath et al. 2007; Niwa et al. 2009; Stavridis et al. 2007). We and others found that *Pten*, a negative regulator of AKT, is among the top hits in genetic screens for drivers of ESC differentiation, highlighting the importance of downregulating the PI3K/AKT pathway to ensure timely exit from the naïve state (Betschinger et al. 2013; Lackner et al. 2021; Leeb et al. 2014; Li et al. 2018; Villegas et al. 2019). However, how exactly *Pten* regulates the exit from naïve pluripotency remains elusive.

In this study, we find that FoxO transcription factors are regulated by AKT and play a previously unrecognized but critical role in the transition from naïve to formative pluripotency. Our findings indicate that AKT acts as a gatekeeper by maintaining FoxO TFs in the cytoplasm in the naïve state. However, PTEN reduces AKT signalling at the initiation of differentiation, allowing FoxO TFs to relocalize to the nucleus where they play a pivotal role in facilitating the transition from naïve to formative pluripotency by regulating a switch in operative GRNs. Our findings uncover an intricate mechanism that regulates the orderly transition between gene regulatory networks that maintain distinct pluripotent states.

## RESULTS

### PTEN-mediated downregulation of AKT activity results in timely exit from the naïve pluripotent state

We previously found that mESCs lacking *Pten* exhibit a pronounced defect in the exit from naïve pluripotency (Lackner et al. 2021). Indeed, 24h after 2i-removal in N2B27 medium (N24), *Pten* KO mESCs displayed higher Rex1-GFPd2 (Rex1-GFP) reporter activity than wild-type (WT) cells (*Fig. 1A, B*). *Rex1* is specifically expressed in the naïve state, and its downregulation coincides with irreversible commitment to differentiation (Betschinger et al. 2013; Wray et al. 2011). This defect in exit from the naïve state in *Pten* KO ESCs was rescued by expressing *Pten* through transfection of a plasmid encoding 3xFLAG-PTEN (*Supplementary Fig. 1A, B*).

**Fig. 1.**
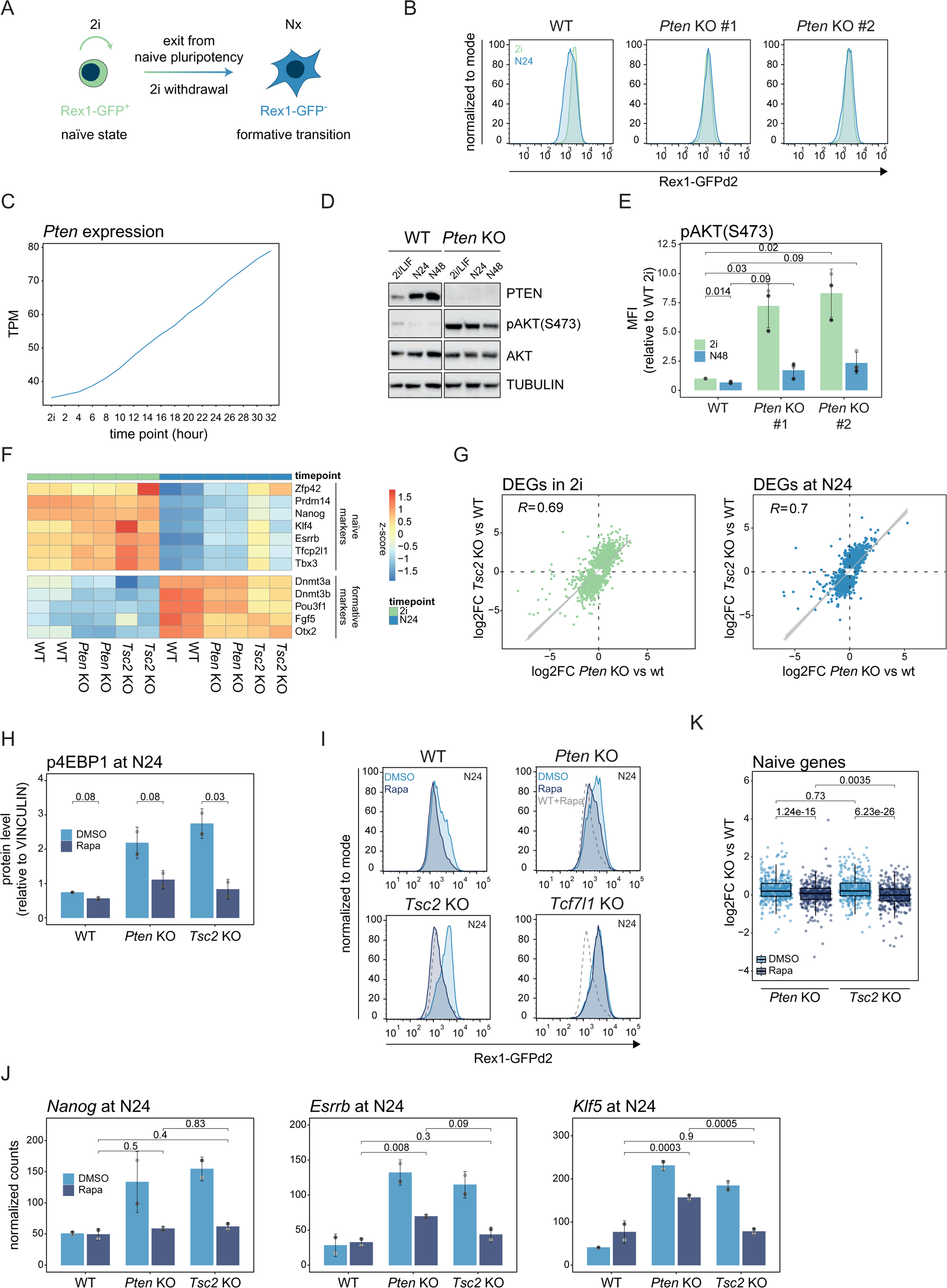
mTORC1 inhibition only partially rescues the differentiation defect of *Pten* KO ESCs. A, Schematic representation of our experimental cell line. Rex1-GFP reporter levels are measured at the indicated time point in N2B27-only medium (Nx) after 2i withdrawal to assess the differentiation state. B, Flow cytometry analysis of Rex1-GFP levels in WT and in *Pten* KO cells (two independent clones) in naïve pluripotency supporting conditions (2i, green profiles) and 24h after 2i removal (N24, light blue profiles). One representative of n >10 independent experiments is shown. C, *Pten* expression levels (transcript per million, TPM) as measured by an RNA-seq experiment from a 2 hour-resolved WT differentiation time course (Lackner et al. 2021). D, Western blot analysis for the indicated proteins in WT and *Pten* KO cells in naïve pluripotency supporting conditions (2i/LIF) and 24h (N24) and 48h (N48) after 2i removal. TUBULIN was used as a loading control. E, Quantification of pAkt (S473) levels measured by flow cytometry in WT and *Pten* KO cells in 2i and at N48. Mean and standard deviation (SD) of mean fluorescence intensity (MFI) values from n=3 independent experiments (distinguished by distinct shades of grey) are shown. Normalised MFI values (see Materials and Methods) are presented relative to WT cells in 2i. The indicated *p-values* shows the result of paired, two-tailed t-tests. F, Heatmap showing row-normalised (z-score) expression values of the indicated naïve and formative markers in WT, *Pten* KO and *Tsc2* KO cells in 2i and at N24. G, Scatter plots showing the correlation between differentially expressed genes (DEGs, *p-adj*. ≤ 0.01, |log2foldchange (log2FC)| ≤ 0.5) in *Pten* KO or *Tsc2* KO in 2i (left) and at N24 (right). Pearson’s correlation coefficient (R) values are indicated in each plot. H, Quantification of p4EBP1 protein expression measured by Western blot analysis in WT, *Pten* KO and *Tsc2* KO cells at N24 after treatment with DMSO (light blue) or 20 nM Rapamycin (dark blue). Expression was normalised to VINCULIN. Mean and SD for n=2 independent experiments (distinguished by distinct shades of grey) are shown. Indicated *p-values* show results of paired, two-tailed t-tests. I, Flow cytometry analysis of Rex1-GFP levels in indicated cell lines at N24, after treatment with DMSO (light blue profiles) or with 20 nM Rapamycin (Rapa, dark blue profiles). Rapamycin-treated WT cells are shown as a grey dashed line. One representative of n=5 independent experiments is shown. J, Expression levels (normalised counts) of *Nanog*, *Esrrb* and *Klf5* measured in the indicated cell lines at N24 after treatment with DMSO or 20 nM Rapamycin measured by RNA-seq. Mean and SD of n=2 independent experiments (distinguished by distinct shades of grey) are shown. *p-adj. values* resulting from DESeq analysis between pairwise comparisons of Rapa-treated cell lines are indicated in the plot. K, Box plot showing the expression of naïve genes (combination of naïve early and naïve late genes as defined in Carbognin et al. 2023) in *Pten* and *Tsc2* KO cells treated with DMSO or Rapamycin as measured by RNA-seq. Data is shown as log2FC relative to WT. The resulting *p-values* from two-tailed Wilcoxon signed rank tests are indicated in the plot.

*Pten* mRNA and protein levels increase during exit from naïve pluripotency, while phospho-AKT (pAKT) levels are concomitantly reduced (*Fig. 1C-E*), suggesting that PTEN may promote the transition to formative pluripotency by decreasing AKT activity. In *Pten* KOs, levels of phospho-AKT were significantly higher than in WT cells (*Fig. 1D*). Thus, we set out to delineate the molecular mechanism by which PTEN-mediated AKT inhibition might drive exit from naïve pluripotency.

When active, AKT phosphorylates several targets that play crucial roles in distinct cellular processes. Among AKT targets, TSC2 (Tuberous Sclerosis 2), GSK3 (Glycogen Synthase Kinase 3), and the FOXO (Forkhead box O) class of transcription factors are strong candidates for mediating changes in mESC differentiation states. Indeed, these factors have been identified as hits in genetic screens for differentiation drivers (Betschinger et al. 2013; Lackner et al. 2021; Leeb et al. 2014; Li et al. 2018; Villegas et al. 2019), suggesting that the AKT/mTORC1, AKT/GSK3 and AKT/FoxO signalling axes could all participate in the regulation of the transition from naïve to formative pluripotency.

Therefore, we started by investigating the involvement of AKT/mTORC1. AKT-mediated phosphorylation leads to TSC2 inhibition and consequent mTORC1 activation. mTORC1 is one of two distinct complexes containing the serine/threonine protein kinase mTOR. mTORC1 regulates essential cellular processes including cell growth, protein synthesis and autophagy via phosphorylation of S6K, 4EBP-1 and ULK1, respectively (Yu and Cui 2016). ESCs lacking *Tsc2* retained higher Rex1-GFP and NANOG levels at N24 compared to WT cells, similar to *Pten* KO cells (Lackner et al. 2021) (*Supplementary Fig. 1C, D*).

To evaluate whether *Pten* acts through the AKT/mTORC1 axis during exit from naïve pluripotency, we inspected RNA sequencing (RNA-seq) data from *Pten* and *Tsc2* KO mESCs (Lackner et al. 2021) (*Supplementary Fig. 1E*). Both KOs showed delayed downregulation of naïve and delayed upregulation of formative marker genes at N24, with more pronounced effects observed in *Tsc2* KOs (*Fig. 1F*). Both in 2i and at N24, *Pten* and *Tsc2* KOs deregulated a similar set of genes (differentially expressed genes, DEGs; *Fig. 1G*). In 2i, DEGs were enriched for terms associated with lysosomal and metabolic regulation, in line with the known role of mTORC1 in regulating those processes (Betschinger et al. 2013; Villegas et al. 2019; Yu and Cui 2016) (*Supplementary Fig. 1F*). Consistent with the observed naïve exit defect, genes upregulated at N24 were enriched for terms related to pluripotency (*Supplementary Fig. 1G*).

Next, we inspected the phosphorylation level of direct targets of mTORC1 in WT, *Pten* KO and *Tsc2* KO ESCs. Phospho-4EBP1 (p4EBP1) and phospho-S6K (pS6K) were similarly increased in both KOs, confirming that *Pten* or *Tsc2* deletion increases mTORC1 pathway activity. Addition of the mTORC1 inhibitor Rapamycin reduced mTORC1 activity in both KOs (*Fig. 1H and Supplementary Fig. 1H, I*).

We then assessed whether this reduction restored normal differentiation potential. Rapamycin treatment promoted faster downregulation of Rex1-GFP in WT ESCs, and resulted in a complete rescue of the differentiation defect of *Tsc2* KO cells (*Fig. 1I*), in line with previously published data (Betschinger et al. 2013). However, in *Pten* KO cells, Rapamycin achieved only a partial rescue. Restoration of differentiation potential was specific for ESCs depleted for members of the PI3K/AKT/mTOR pathway, and differentiation-defective ESCs lacking the WNT/GSK3 pathway effector *Tcf7l1* (Lackner et al. 2021; Wray et al. 2011) did not restore Rex1-GFP downregulation kinetics upon Rapamycin treatment (*Fig. 1I*).

To further evaluate the extent of phenotypic rescue of *Pten* KO cells by Rapamycin, we performed RNA-Seq on WT, *Pten* KO and *Tsc2* KO mESCs, differentiated in presence of DMSO or Rapamycin (*Supplementary Fig. 1J*). We observed TF-specific changes in naïve pluripotency expression upon Rapamycin treatment. While addition of Rapamycin restored *Nanog* expression to WT levels in both *Pten* and *Tsc2* KOs, *Esrrb* and *Klf5* expression remained high in *Pten* KOs. Furthermore, a set of 436 previously identified naïve pluripotency-specific genes (Carbognin et al. 2023) consistently showed a significantly stronger reduction in expression levels upon Rapamycin treatment in *Tsc2* compared to *Pten* KOs (*Fig. 1K*). Together, these results show that the *Pten* KO phenotype is not exclusively determined by hyperactivity of mTORC1.

This prompted us to first investigate the role of AKT-mediated phosphorylation of GSK3 on Serine 9, which tags it for degradation and thereby stabilises ꞵ-catenin (Wray et al. 2011). GSK3 phosphorylation was recently proposed to be crucial for maintaining pluripotency in *Pten* KO mESCs (Wang et al. 2020). Indeed, phospho-GSK3 (pGSK3) levels are increased in *Pten* KO cells (*Supplementary Fig. 1K*). We hypothesised that if indeed deactivation of the GSK3-TCF7L1 axis of the WNT pathway leads to the differentiation defect in *Pten* KO cells, then the resulting WNT pathway hyperactivity should be epistatic to the pharmacological inhibition of GSK3 by CHIRON. Such an epistatic interaction was observed in *Tcf7l1* KOs, where the activity of the ꞵ-catenin destruction complex is rendered obsolete and, hence, the addition of CHIRON had no additional effect (*Supplementary Fig. 1L*). In contrast, treatment of *Pten* KOs with CHIRON resulted in delayed differentiation speeds akin to WT cells. Furthermore, we did not previously observe a transcriptional signature typical of increased WNT activity in *Pten* KO ESCs (Lackner et al. 2021). This suggests that AKT-hyperactivity-dependent phosphorylation of GSK3 in *Pten* mutants has little direct impact on the exit from naïve pluripotency.

Collectively, these data show that *Pten* regulates pathways, in addition to mTORC1, relevant for proper exit from naïve pluripotency. As the role of GSK3 appeared minor, we focussed our attention on FoxO TFs.

### AKT-dependent nuclear translocation of FoxO transcription factors promotes the exit from naïve pluripotency

FoxO TFs regulate several crucial cellular processes, including cell cycle, apoptosis and DNA repair (Herman, Todeschini, and Veitia 2020). AKT-mediated phosphorylation retains FoxO TFs in the cytoplasm. In WT and *Pten* KO ESCs cultured in 2i, immunofluorescence (IF) analysis revealed a largely cytoplasmic localization of FOXO1. Upon exit from the naïve state, we observed that FOXO1 translocated to the nucleus, concomitant with reduced ESRRB levels (*Fig. 2A, B*) and a downregulation of AKT activity (*Fig. 1D, E*) (Yang et al. 2019). IF experiments revealed that increased nuclear FOXO1 levels could be observed as early as 8h after 2i withdrawal (N8), and increased further until N24. Nuclear translocation of FOXO1 at the onset of ESC differentiation was severely impaired in *Pten* KO cells, which maintained a clear cytoplasmic localization at N24 (*Fig. 2A, C*). Nucleo-cytoplasmic fractionation experiments showed similar results for FoxO3 (*Supplementary Fig. 2A*). Supporting a functional relevance of nuclear shuttling of FoxO TFs *in vivo*, we detected nuclear FoxO1 in the OTX2 positive epiblast of E4.75 and E5.5 embryos (*Fig. 2D*). In contrast, extraembryonic tissues showed lower FoxO1 levels and a more pronounced cytoplasmic FoxO1 localization.

**Fig. 2.**
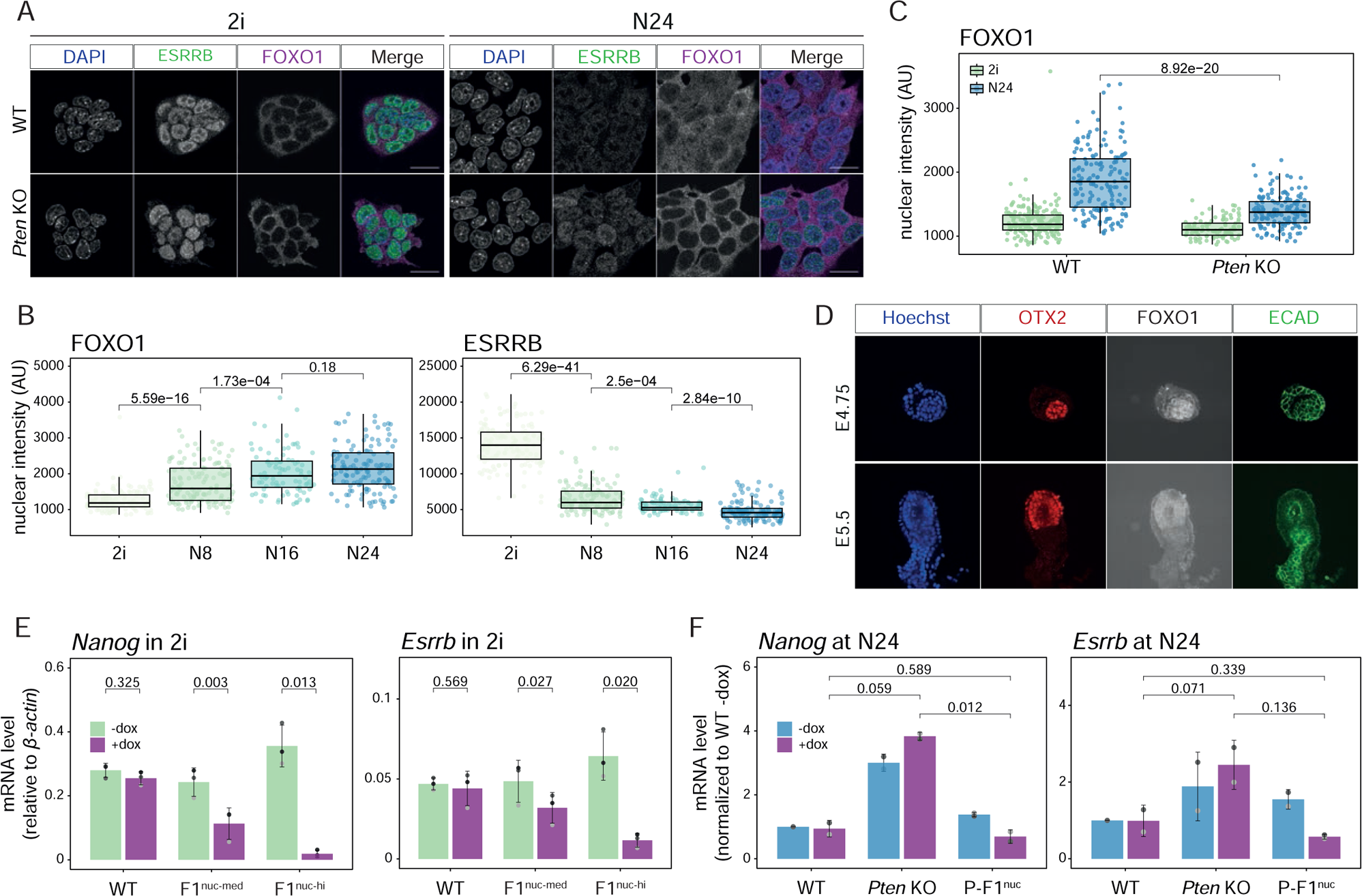
FoxO TFs translocate into the nucleus upon exit from naive pluripotency. A, Confocal microscopy images after IF showing FOXO1 (purple) and ESRRB (green) in WT and *Pten* KOs in 2i and at N24. DAPI staining is shown in blue. One representative of n=2 independent experiments is shown. Scale bar = 20 μM. B, Quantification of FOXO1 and ESRRB nuclear intensity (arbitrary units, AU) measured from confocal images of WT cells in 2i (n=135) and 8h (N8, n=136), 16h (N16, n=88) and 24h (N24, n=127) after 2i withdrawal. Data from n=2 independent experiments is shown. The indicated *p-value* shows the result of a two-tailed Wilcoxon rank sum test. C, Quantification of FOXO1 nuclear intensity (in arbitrary unit, AU) measured from confocal images of WT (n=189 in 2i and n=164 at N24) and *Pten* KO cells (n=100 in 2i and n=153 at N24) as in A. Data from n=2 independent experiments is shown. The indicated *p-values* show the results of two-tailed Wilcoxon rank sum tests. D, Confocal microscopy images after IF showing FOXO1 (white), OTX2 (red) and ECAD (green) in WT E4.75 and E5.5 embryos. Hoechst staining is shown in blue. One representative of n=2 independent experiments is shown. E, Expression levels of *Nanog* and *Esrrb* measured by RT-qPCR in WT cells expressing 3xFLAG-FoxO1^nuc^ (FoxO1^nuc-med^ and FoxO1^nuc-hi^) in 2i after 8 hours treatment with 500 ng/ml doxycycline (+dox, purple) and in untreated controls (-dox, green). Mean and SD of n=3 independent experiments (distinguished by distinct shades of grey) are shown. Expression was normalised to *β-actin*. *p-values* show results of paired, two-tailed t-tests. F, Expression levels of *Nanog* and *Esrrb* measured by RT-qPCR in *Pten* KO cells expressing 3xFLAG-FoxO1^nuc^ (P-FoxO1^nuc^) at N24 after 8 hours treatment with 500 ng/ml doxycycline (+dox, purple) and in untreated controls (-dox, light blue). Control WT cells and *Pten* KO cells are also included. Mean and SD of n=2 independent experiments (distinguished by distinct shades of grey) are shown. Expression was normalised to *β-actin* and shown as relative to WT in -dox condition. *p-values* show results of paired, two-tailed t-tests.

To test whether nuclear translocation of FOXO1 was indeed AKT-dependent, we analysed the nuclear vs. cytoplasmic localization of FOXO1 after treatment with the specific allosteric AKT inhibitor MK-2206 (Hirai et al. 2010). MK-2206 treatment elevated levels of nuclear FOXO1, in both 2i and at N24, and resulted in expedited downregulation of Rex1-GFP in WT cells (*Supplementary Fig. 2B-D*), consistent with a requirement for reduced AKT activity to enable and possibly trigger the exit from naïve pluripotency.

Together, our results supported the hypothesis that impaired nuclear translocation of FoxO TFs at the onset of differentiation contributes to the inability of *Pten* KO ESCs to properly exit the naïve state. To further test this hypothesis, we used doxycycline-induced expression of a constitutively nuclear version of FoxO1 (3xFLAG-FoxO1^nuc^) (Nakae et al. 2001). This treatment was sufficient to extinguish naïve pluripotency in WT cells cultured in 2i within 8h hours in a dose-dependent manner (*Fig. 2E*, *Supplementary Fig. 2E-G*). Furthermore, 3xFLAG-FoxO1^nuc^ expression in *Pten* KO ESCs rescued their differentiation defect (*Fig. 2F*, *Supplementary Fig. 2H, I*). We noticed that prolonged exposure to high levels of nuclear FoxO1 (>24h) was cytotoxic, probably due to induction of the pro-apoptotic programme by FoxO1 (Greer and Brunet 2005).

In summary, our data show that AKT-mediated nuclear translocation of FoxO1, and possibly other FoxO TFs, is essential for the timely exit from naïve pluripotency. We conclude that lack of FoxO TF nuclear translocation underlies, at least in part, the differentiation defect observed in *Pten* KO mESCs.

### FoxO TFs bind to enhancers that are activated during the naïve to formative transition

To explore the role of FoxO TFs in the transition from naïve to formative pluripotency, we performed chromatin immunoprecipitation followed by next-generation sequencing (ChIP-seq) analysis for FOXO1 and FOXO3 in WT and *Pten* KO cells. This analysis was performed in 2i and at N24, that is, 24 hours into the formative transition (*Fig. 3A and Supplementary Fig. 3A-C*). In WT cells, we observed a strong increase in the number of genomic loci bound by FOXO1 and FOXO3 at N24 compared to 2i. We detected 1314 and 391 peaks in 2i and 2840 and 623 peaks at N24 for FoxO1 and FoxO3, respectively. This is consistent with increased nuclear localization of FoxO TFs at the exit from naïve pluripotency. In agreement with the largely cytoplasmic localization in the absence of *Pten*, FOXO-TF ChIP-seq signals were barely detectable in *Pten* KO cells (*Supplementary Fig. 3B, C*). Of note, due to low amounts of precipitated DNA in 2i, ChIP-seq library-prep will most likely have amplified signal in 2i relative to N24; hence a direct quantitative comparison between 2i and N24 FOXO-TF signal is not possible.

**Fig. 3.**
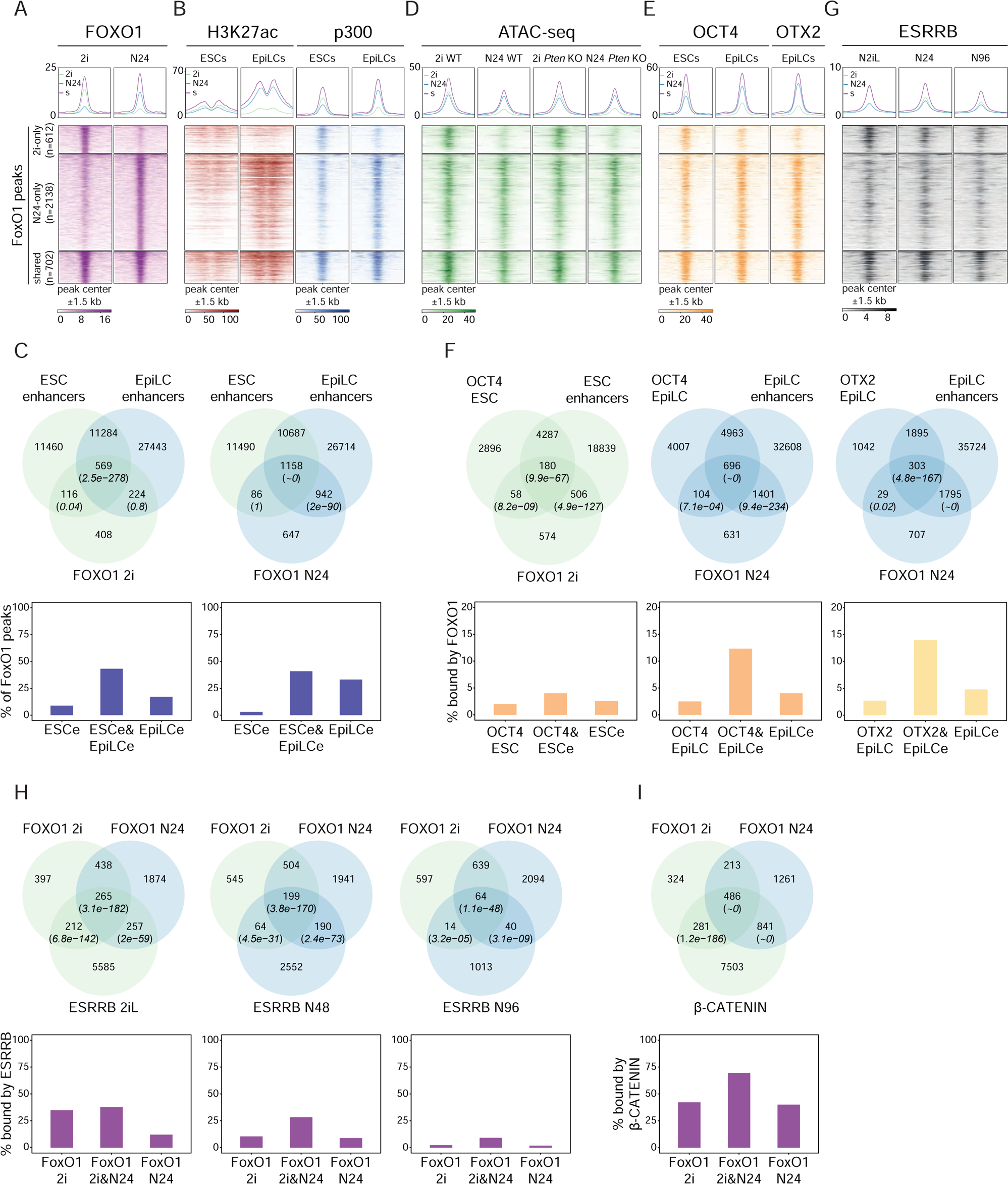
Chromatin dynamics of FoxO TFs upon exit from the naïve pluripotent state. A, Heatmap showing FOXO1 signal in a 1.5 kb window around the center of FoxO1 peaks, divided in 2i-only peaks (n=612, green), N24-only peaks (n=2138, light blue) and shared peaks (n=702, purple), in WT cells in 2i or at N24. B, Heatmaps showing H3K27ac (left) and p300 (right) signals in ESCs or EpiLCs (from Buecker et al. 2014) in a 1.5 kb window around the center of FoxO1 peak categories as defined in A. C, Venn diagrams showing the overlap between FoxO1 peaks in 2i or at N24 with ESC and EpiLC enhancers (*top*). Barplots showing the % of FoxO1 peaks that overlap with ESC-specific (ESCe), EpiLC-specific (EpiLCe) or shared (ESCe&EpiLCe) enhancers, (*bottom*). The resulting *p-values* from hypergeometric tests of the overlaps are shown. D, Heatmap showing ATAC-seq signal in a 1.5 kb window around the center of FoxO1 peak categories as defined in A in WT and *Pten* KO cells in 2i or at N24. E, Heatmaps showing OCT4 (left) and OTX2 (right) signals in ESCs or EpiLCs (from Buecker et al. 2014) in a 1.5 kb window around the center of FoxO1 peak categories as defined in A. F, Venn diagrams showing the overlap between FoxO1 peaks in 2i or at N24 with Oct4 or Otx2 peaks in ESCs or EpiLCs (*top*). Barplots showing the % of OCT4- or OTX2-bound enhancers that overlap with FoxO1 peaks (*bottom*). The resulting *p-values* from hypergeometric tests of the overlaps are shown. G Heatmap showing ESRRB signal in WT cells in 2iL, N48 and N96 (from Carbognin et al. 2023) in a 1.5 kb window around the center of FoxO1 peak categories as defined in A. H, Venn diagrams showing the overlap between FoxO1 peaks in 2i and at N24 with Esrrb peaks in 2iL, N48 or N96 (*top*). Barplots showing the % of FoxO1 peaks (2i-only, N24-only or shared) that overlap with Esrrb peaks (*bottom*). The resulting *p-values* from hypergeometric tests of the overlaps are shown. I, Venn diagrams showing the overlap between FoxO1 peaks in 2i and at N24 with β-catenin peaks in 2iL, N48 or N96 (*top*). Barplots showing the % of FoxO1 peaks (2i-only, N24-only or shared) that overlap with β-catenin peaks (*bottom*). The resulting *p-values* from hypergeometric tests of the overlaps are shown.

FOXO1 and FOXO3 ChIP-seq data showed a significant overlap. Almost 50% of FOXO3 peaks overlapped with FOXO1 peaks, and FOXO1 signal was detected at virtually all FoxO3 peaks (*Supplementary Fig. 3C, D*). We divided FOXO1- and FOXO3-bound regions into three groups depending on peak-calling results: 2i-only peaks (612 for FOXO1 and 152 for FOXO3), N24-only peaks (2138 for FOXO1 and 384 for FOXO3) and shared peaks (702 for FOXO1 and 239 for FOXO3) (*Fig. 3A*). All peak categories were enriched for FoxO motifs (*Supplementary Table 1*). Altogether, this supports the specificity of our ChIP-seq analysis.

FoxO TF peaks were located mainly outside of promoter regions (*Supplementary Fig. 3E*), indicating a potential contribution of FoxO TFs to enhancer regulation. To test this, we utilised published ChIP-seq datasets (Buecker et al. 2014) for the enhancer marks H3K27ac and p300, obtained in ESCs and in EpiLCs. EpiLCs represent the *in vitro* counterpart of formative epiblast cells of the E5.5 blastocyst (Hayashi et al. 2011) and correspond to a developmental state similar to our cells at N24. We found that strong H3K27ac and p300 signals in EpiLCs overlapped with the regions we identified as bound by FoxO TFs at N24 (*Fig. 3B*). Moreover, regions bound by FoxO TFs in 2i were significantly enriched in ESC-specific enhancers and enhancers active in ESCs and EpiLCs (Thomas et al. 2021) (*Fig. 3C*). EpiLC-specific enhancers were not significantly bound by FOXO1 in 2i. In contrast, N24 FoxO1-peaks significantly overlapped with shared and EpiLCs-specific enhancers while showing much less overlap with ESC-specific enhancers. We found that a total of xX% of FoxO1 peaks at N24 overlaps with EpiLCs enhancers. This points towards a role for FoxO TFs in activating formative-specific regulatory regions upon exit from the naïve state. Consistently, we detected a significant enrichment of FoxO1 and FoxO3 motifs in EpiLC-specific enhancers (*Supplementary Fig. 3F*).

We then performed ATAC-seq in WT and *Pten* KO cells, in 2i and at N24 (*Fig. 3D, Supplementary Fig. 3G*). Regions that showed FoxO-TF binding exclusively in 2i were open only in the naïve state and showed weak ATAC-seq signal at N24. However, regions exhibiting FoxO TF binding at N24 showed near equal ATAC-seq signal in both 2i and at N24. At the resolution of our analysis, ATAC-seq signal was indistinguishable between WT and *Pten* KO cells, in which FoxO-TF translocation to the nucleus is largely abolished. This suggests that FoxO TFs are not likely to act as pioneer factors that open up chromatin during the naïve to formative transition. Instead, our data is consistent with a role of FoxO TFs in activating already open poised chromatin (Eijkelenboom et al. 2013) during the exit from naïve pluripotency.

OCT4 and OTX2 are key regulators of general and formative pluripotency, respectively. OTX2 causes relocation of OCT4 from ESC- to EpiLC-specific enhancers upon exit from the naïve state (Buecker et al. 2014). We found that regions bound by FoxO1 at N24 showed a significant and strong overlap with OCT4-bound regions detected in EpiLCs. This contrasted with lower overlap between regions bound by FoxO1 in 2i and OCT4 in ESCs (*Fig. 3E, F*). A large portion of 14% of OTX2-bound EpiLC enhancers were also bound by FoxO1 at N24. In addition, our analysis clearly showed that colocalization of FOXO1 with OCT4 and OTX2 occurs nearly exclusively on enhancers, suggesting that FoxO TFs regulate enhancer activation in cooperation with pluripotency-state specific TFs.

Recently it was shown that *Esrrb,* originally identified as a naïve pluripotency-specific TF, performs additional functions in the initiation of the formative transcription programme (Carbognin et al. 2023). To investigate whether FoxO-TFs cooperate with ESRRB, we examined the overlap on chromatin between FOXO1 and ESRRB throughout the pluripotency continuum (*Fig. 3G, H and Supplementary Fig. 3H, I*). These analyses showed that 35% of 2i-specific and 38% of shared FoxO1 peaks are also bound by ESRRB in 2i (*Fig. 3H*). Regions bound by ESRRB at N48 still showed a highly significant overlap with FoxO1 binding, with 28% of shared FoxO1 peaks also decorated by ESRRB at N48. These loci were in close proximity to, and thus potentially regulate, known formative (*Otx2*, *Dnmt3a*, *Lef1*) and naive marker genes (*Nanog*, *Esrrb*, *Zfp42*, *Klf2*, *Klf4*, *Klf5*, *Tfcp2l1*, *Nr5a2*, *Prdm14*). Overall, regions co-bound by FOXO1 and ESRRB showed the strongest signals for enhancer marks, indicating that the concomitant presence of FoxO TFs and ESRRB triggers the strongest transcriptional response (*Supplementary Fig. 3H*, I).

FoxO TFs have been shown to interact with the WNT pathway effector β-catenin to enhance transcriptional output (Essers et al. 2005). In line with this, a highly significant 50% of all FoxO1 bound regions were also occupied by β-catenin (*Fig. 3I*), suggesting a functional interaction of FoxO TFs and β-catenin also during the transition to formative pluripotency.

Our findings reveal that FoxO TFs bind to enhancers that are activated upon the exit from naïve pluripotency. This suggests that FoxO TFs cooperate with core components of the naïve and formative TF-repertoire to ensure faithful firing of the formative GRN.

### FoxO TFs instruct the rewiring of the naïve to the formative GRN

We next sought to identify the transcriptional consequences of FoxO-TF induced changes to chromatin at the exit from naïve pluripotency. We specifically wanted to know which components of the formative state specific GRN might be functionally dependent on regulation by FoxO TFs. To this end, we compared the changes in transcript levels upon exit from naïve pluripotency (Lackner et al. 2021) between distinct sets of FoxO1-bound genes as defined above. Overall, FoxO1 targets were highly enriched in genes differentially expressed within the first 24h of ESC differentiation (*Supplementary Fig. 4A*). 2i-specific FoxO1 targets were overall downregulated at N24, while N24-specific targets showed an overall upregulation during naïve exit (*Fig. 4A, Supplementary Fig. 4B*). In contrast, the expression of targets bound in both conditions did not show a consistent directional change in gene expression during the exit from naïve pluripotency. To further investigate the exact expression kinetics of FoxO1 target genes during the exit from naïve pluripotency, we followed their transcript levels across a 32h differentiation time course at a 2h-resolution (Lackner et al. 2021). We found that 2i-specific FoxO1 targets showed a largely continuous downregulation, N24-specific targets a continuous upregulation, and 2i&N24 targets a lack of clear directionality (*Supplementary Fig. 4C*).

**Fig. 4.**
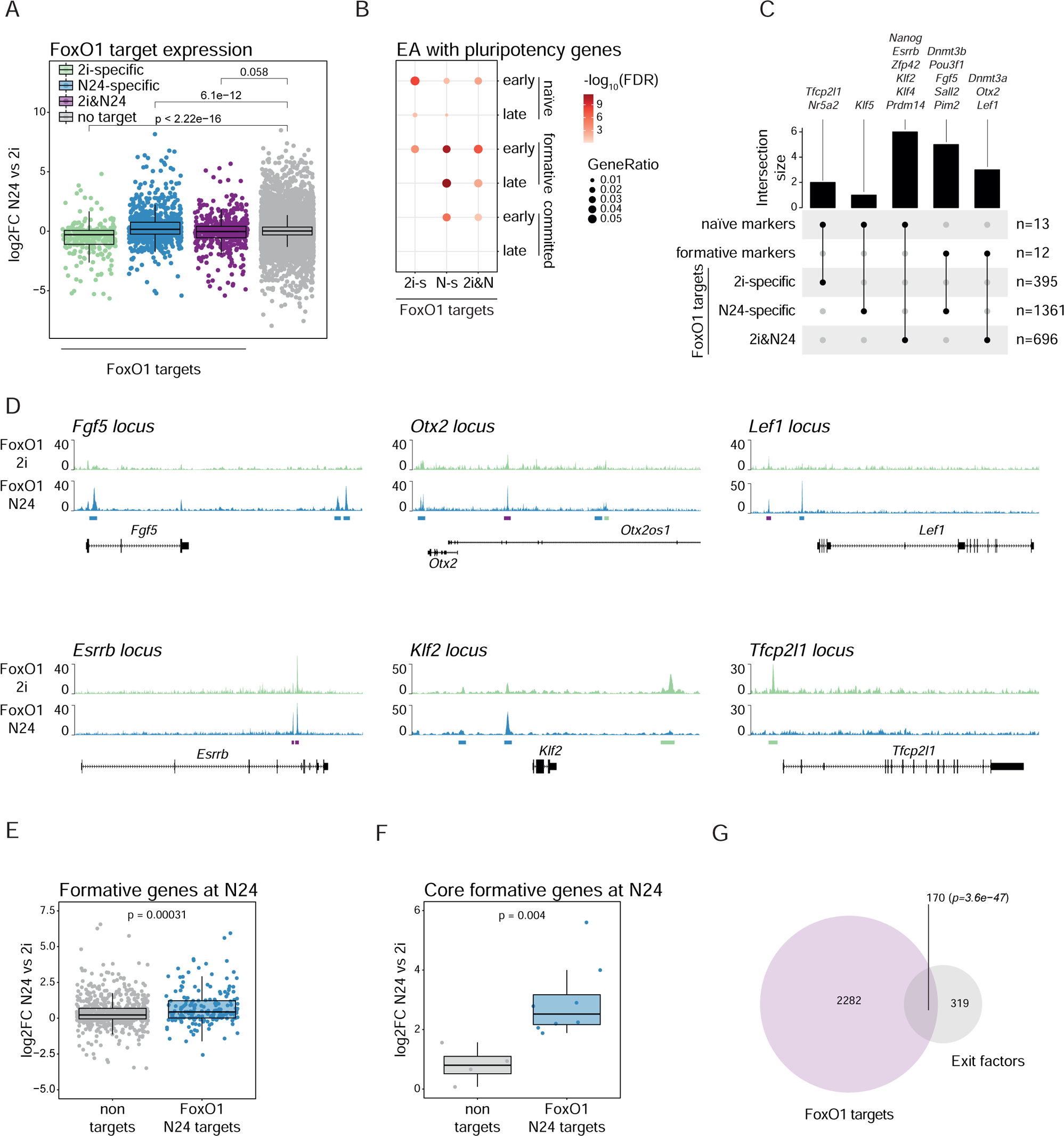
FoxO TF targets are key players in the naïve to formative pluripotency transition. A, Box plot showing the expression of 2i-specific (green), N24-specific (light blue) and 2i&N24 (purple) FoxO1 targets in WT cells at N24 as measured by RNA-seq. Data is shown as log2FC relative to 2i. The resulting *p-values* from two-tailed Wilcoxon rank sum tests are indicated in the plot. B, Enrichment analysis (EA) of FoxO1 targets within lists of naïve, formative and committed genes as defined in Carbognin et al. 2023. Dot colour and size indicate the FDR-values (only FDR ≤ 0.05 are shown) and the GeneRatio (ratio between the overlap size and the category size), respectively. C, Upset plot showing the overlap of FoxO1 targets with core naïve and formative marker genes. D, Genome browser snapshots showing FoxO1 ChIP signal in WT cells over selected formative (top) and naïve (bottom) core marker genes in 2i (green) and at N24 (light blue). Called peaks are also indicated in the plots (2i-only peaks: green, N24-only peaks: light blue, shared peaks: purple). E, Box plot showing the expression of formative genes (combination of formative early and formative late, as defined in Carbognin et al. 2023) in WT cells at N24, divided into FoxO1 N24 targets (light blue) or non-targets (grey), as measured by RNA-seq. Data is shown as log2FC relative to 2i. The resulting *p-value* from two-tailed Wilcoxon rank sum tests is indicated in the plot. F, Box plot showing the expression of core formative genes (as defined in the text) in WT cells at N24, divided into FoxO1 N24 targets (light blue) or non-targets (grey), as measured by RNA-seq. Data is shown as log2FC relative to 2i. The resulting *p-value* from two-tailed Wilcoxon rank sum test is indicated in the plot. G, Venn diagram showing the overlap between FoxO1 targets (purple) and exit factors (Lackner et al. 2021) (grey). The resulting *p-value* from a hypergeometric test of the overlap is shown.

Recent work reported transcriptome analysis of differentiating mESCs up to 96 hours after 2i withdrawal and defined 6 distinct groups based on gene expression kinetics: naïve early and naïve late (downregulated early or late upon naïve exit), formative early and formative late (upregulated during naïve exit), committed early and committed late (upregulated late during naïve exit) (Carbognin et al. 2023). We found that 2i-specific FoxO1 targets were enriched for naïve and formative early genes, whereas N24-specific and 2i&N24 FoxO1 targets were enriched for naïve early, formative (early and late) and committed early genes (*Fig. 4B*).

As this result indicated a link between FoxO TFs and central components of the naïve and formative GRNs, we further tested whether FoxO1 binds and potentially regulates core members of the naïve or formative GRN. We found that 9 out of 13 core naïve marker genes (modified from Kalkan et al. 2019) were bound by FoxO1 in 2i (*Tfcp2l1*, *Nr5a2*) or in both 2i and at N24 (*Nanog*, *Esrrb*, *Zfp42*, *Klf2*, *Klf4, Prdm14*). Conversely, 8 out of 12 key formative marker genes (hereafter referred to as core formative genes) (Kalkan et al. 2019) were bound by FoxO1 at N24 (*Dnmt3b*, *Pou3f1*, *Fgf5*, *Sall2*, *Pim2*) or under both conditions (*Dnmt3a*, *Otx2*, *Lef1*) (*Fig. 4C, D and Supplementary Fig. 4D, Supplementary Table 2*). Notably, formative genes (early and late, as described above) and core formative genes that are direct FoxO1 targets showed a significantly stronger upregulation during the exit from naïve pluripotency than non FoxO-targets (*Fig. 4E, F*).

Among the FoxO1 targets, we identified a significant number of genes that were also found in a genetic screen for genes controlling the exit from naïve pluripotency, here referred to as “exit factors” (*Fig. 4G*) (Lackner et al. 2021; Leeb et al. 2014). Most exit factors show little dynamic change in gene expression during the exit from naïve pluripotency. However, among those that are upregulated during naïve exit, FoxO1 targets showed significantly stronger regulation (*Supplementary Fig. 4E*). This suggests that FoxO TFs function as upstream regulators of the cell fate transition from naïve to formative pluripotency by facilitating the activity of multiple processes required for proper differentiation.

Further suggesting a causal link between inactivation of the FoxO-signalling axis in *Pten* KOs and the differentiation delay in these mutants, 22% of genes differentially expressed between WT and *Pten* KOs at N24 were FoxO1 target genes (*Supplementary Fig. 4F*). Furthermore, those components of the core formative GRN that are FoxO1 targets exhibited a stronger deficiency in upregulation compared to non-targets in *Pten* mutant cells at N24 (*Supplementary Fig. 4G*).

In sum, our data argue that FoxO1 is a key regulator of the naïve to formative transition by binding to and regulating major components of the naïve and formative specific GRNs. This function is potentially performed in cooperation with TFs such as OCT4, ESRRB, OTX2 and β-CATENIN that are known to regulate multiple pluripotency states and transitions.

### FoxO TFs are required for the exit from naïve pluripotency

To investigate a potential requirement for FoxO1 for the transition from naïve to formative pluripotency, we analysed the differentiation capacity of cells after FoxO1 depletion. To this end, we turned to knockdown (KD) experiments using short-interfering RNAs (siRNAs). Treatment with siRNAs against *Foxo1* resulted in the downregulation of transcript and protein levels (*Supplementary Fig. 5A, B*). Cells with reduced levels of *Foxo1* (siFoxO1) retained higher expression of Rex1-GFP at N24 compared to cells treated with scrambled siRNAs (siScr), suggesting that *Foxo1* is required for proper exit from naïve pluripotency (*Fig. 5A*).

**Fig. 5.**
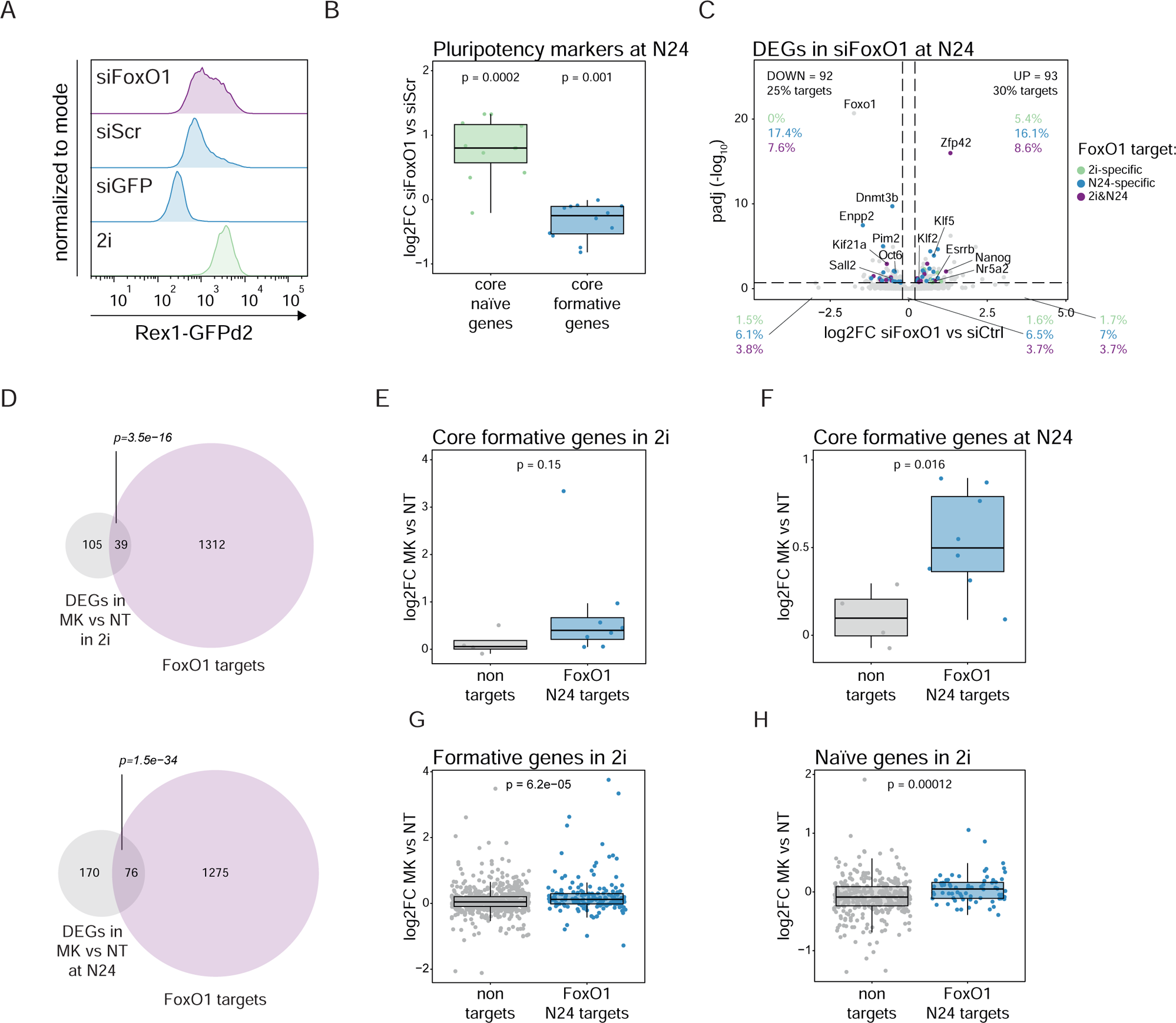
Interference with FoxO1 nuclear shuttling impairs the transition from the naïve to the formative GRN. A, Flow cytometry analysis of Rex1-GFP levels in WT cells transfected with control (siGFP and siScr) or siRNAs targeting FoxO1 (siFoxO1). One representative of n=3 independent experiments is shown. B, Box plot showing the expression of naïve (green) and formative (blue) core genes in siFoxO1 at N24 as measured by RNA-seq. Data is shown as log2FC relative to siCtrl. *p-values* from two-tailed Wilcoxon signed rank tests are indicated in the plot. C, Volcano plot showing RNA-seq data of WT cells at N24, treated with siRNA against *Foxo1*. DEGs (*p-adj.* ≤ 0.2) that are bound by FoxO1 are colour coded depending on whether they are bound only in 2i (green), only at N24 (light blue) or in both conditions (purple). Selected naïve and formative genes are indicated in the plot. For each quadrant, percentages (%) of FoxO1 2i-only, N24-only and 2i&N24 targets are indicated. D, Venn diagrams showing the overlap between FoxO1 targets (purple) and genes differentially expressed upon MK-2206 treatment in 2i (left) or at N24 (right) (DEGs, *p-value* ≤ 0.05, grey). The resulting *p-values* from hypergeometric tests of the overlaps are shown. E, Box plot showing the expression of core formative genes in WT cells after MK-2206 treatment in 2i, divided into FoxO1 N24 targets (light blue) or non-targets (grey). Data is shown as log2FC relative to NT cells. The resulting *p-value* from two-tailed Wilcoxon rank sum test is indicated in the plot. F, Box plot showing the expression of core formative genes in WT cells after MK-2206 treatment at N24, divided into FoxO1 N24 targets (light blue) or non-targets (grey). Data is shown as log2FC relative to NT cells. The resulting *p-value* from two-tailed Wilcoxon rank sum test is indicated in the plot. G, Box plot showing the expression of formative genes (combination of formative early and formative late, as defined in Carbognin et al. 2023) in WT cells treated with the MK-2206 inhibitor in 2i, divided into FoxO1 N24 targets (light blue) or non-targets (grey). Data is shown as log2FC relative to NT cells. The resulting *p-value* from two-tailed Wilcoxon rank sum test is indicated in the plot. H, Box plot showing the expression of naïve genes (combination of naïve early and naïve late, as defined in Carbognin et al. 2023) in WT cells treated with the MK-2206 inhibitor in 2i, divided into FoxO1 N24 targets (light blue) or non-targets (grey). Data is shown as log2FC relative to NT cells. The resulting *p-value* from two-tailed Wilcoxon rank sum test is indicated in the plot.

KEGG pathway enrichment analysis after RNA-seq analysis of WT cells at N24 treated with siFoxO1 showed that upregulated genes (UP) were enriched for signalling pathways associated with pluripotency maintenance (*Supplementary Table 1*), suggesting a fortified naïve pluripotent state and confirming the differentiation defect. Further consistent with the differentiation delay upon FoxO1 depletion, we observed that naïve pluripotency markers failed to be down- and formative markers upregulated upon FoxO1 KD (*Fig. 5B*). Both genes that were up- and downregulated upon FoxO1 depletion were highly enriched in FoxO1 ChIP-seq targets (*Fig. 5C and Supplementary Fig. 5C*), with 30% and 25%, respectively, being bound by FOXO1.

In line with the proposition that the *Pten* KO phenotype is caused, in part, by misregulated FoxO-TF localization, genes upregulated or downregulated in *Pten* KO at N24 were also upregulated or downregulated in siFoxO1-treated cells at N24 (*Supplementary Fig. 5D*). More specifically, FoxO1 target genes showed a stronger deregulation in the absence of either *Pten* or *Foxo1* compared to non-targets (*Supplementary Fig. 5E*).

Our results show that reduced expression of FoxO1 delays the exit from naïve pluripotency, suggesting that FoxO TF-activity is required for the regulation of the naïve to formative pluripotency transition.

### Forced nuclear shuttling of FoxO1 through AKT inhibition is sufficient to promote formative GRN activation in 2i

If nuclear shuttling of FoxO-TFs is indeed involved in initiating the formative GRN, then artificial inactivation of AKT in the naïve pluripotent state should lead to nuclear accumulation of FoxO-TFs and, thus, trigger expression of formative specific genes. To test this proposition, we again employed the allosteric AKT inhibitor MK-2206. MK-2206 treatment leads to a substantial increase in nuclear FoxO1 and a concomitant reduction of Rex1-GFP expression levels, as shown before. To evaluate the global impact of AKT-inhibition in naïve ESCs and during the exit from naïve pluripotency, we performed RNA-seq after MK-2206 treatment in 2i and at N24. In general, MK treatment appeared to push cells towards a more differentiated state, both when added to naïve ESCs or to cells exiting naïve pluripotency (*Supplementary Fig. 5F*). Importantly, putative FoxO1 targets identified by ChIP represented 27% and 30% of all deregulated genes upon MK treatment in 2i cells or at N24, respectively (*Fig. 5D*). Overall, core formative FoxO1 targets *Pou3f1*, *Pim2*, *Dnmt3b*, *Lef1*, *Fgf5*, *Otx2* and *Sall2* showed a stronger response to MK-2206 treatment in both 2i and at N24 than non-targets (*Hes6*, *Sox12*, *Sox3*, *Tead2*) (*Fig. 5E, F*). Consistent results were obtained using a larger set of formative-specific genes (*Fig. 5G*). Interestingly, FoxO1-bound components of the naïve GRN also showed a positive response to MK-2206 treatment in 2i medium (*Fig. 5H*). Showing a specific response to FoxO signalling rather than mTORC1 deregulation, treatment with Rapamycin had no specific effect on FoxO1 targets (*Supplementary Fig. 5G*)

In sum, our work uncovers a novel role for FoxO TFs downstream of AKT signalling in actuating the transition from a naïve to a formative pluripotency specific GRN. Our data shows that this is achieved by FoxO-TFs targeting and regulating large parts of the formative and naïve specific GRNs. Hence, FoxO TFs are pivotal factors in mediating the transition from naïve to formative pluripotency.

## DISCUSSION

In this work, we uncovered a mechanism through which PTEN-mediated AKT regulation controls the transition from naïve to formative pluripotency by regulating nuclear FoxO-TF localization. We demonstrate that FoxO TFs play a fundamental role in orchestrating the exit from naïve pluripotency. Their activity is precisely gated by PTEN and released once the exit from naïve pluripotency commences (*Fig. 6*).

**Fig. 6.**
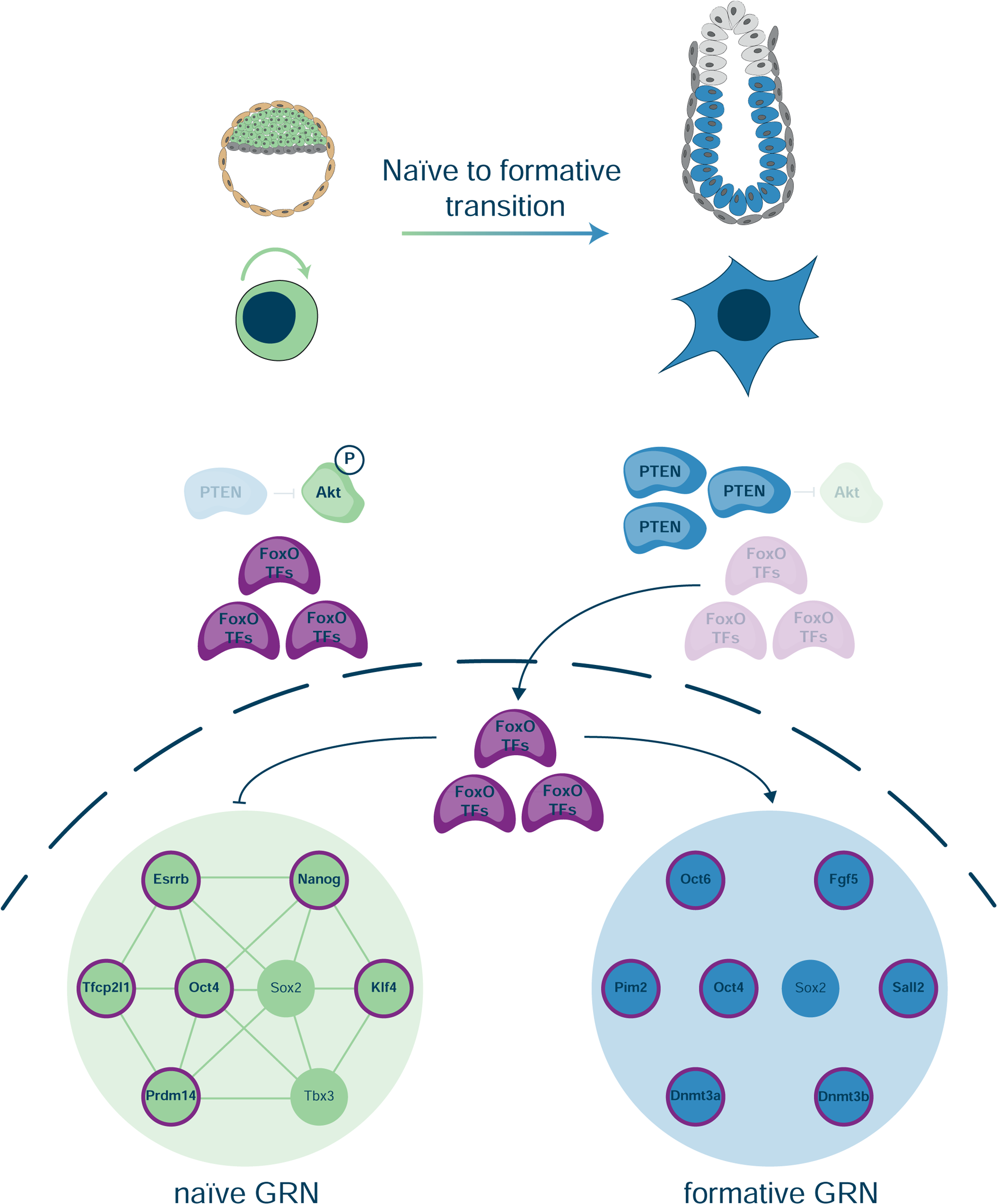
Akt signalling regulation through FoxO TFs of the naïve to formative pluripotency transition. Schematic illustration of the proposed model of FoxO-TF action at the exit from naive pluripotency.

FoxO TFs are well established regulators of multiple fundamental cellular processes including stress response, DNA repair, cell cycle and apoptosis, metabolism and ageing (Carter and Brunet 2007). Although mostly studied for their role in apoptosis, longevity, and cancer, previous studies have reported a function for FoxO TFs in the regulation of cell fate (Greer and Brunet 2005; Paik et al. 2009; Renault et al. 2009; Vilchez et al. 2012, 2013; Webb et al. 2013; Zhang et al. 2011). FoxO TFs are required for the maintenance of neural progenitor cells (NPCs) by preventing neurogenesis *in vivo* and *in vitro* (Paik et al. 2009; Renault et al. 2009; Webb et al. 2013). FoxO4 was proposed to be the only FoxO TF family member necessary for human ESC differentiation into the neuronal lineage (Vilchez et al. 2013), whereas FoxO1 depletion resulted in loss of ESC pluripotency in human and mouse ESCs (Zhang et al. 2011). These data may explain our unsuccessful attempts to generate stable clonal *Foxo1* KO mESCs (data not shown), though the molecular requirement for FoxO1 in ESC self-renewal remains unclear.

Relocating transcription factors from the cytoplasm to the nucleus or *vice versa* allows cells to rapidly respond to changes in signalling inputs by providing a cell state switch with rapid on-off kinetics. A relevant example is the regulation of TFE3, a bHLH transcription factor that is relocated from the nucleus to the cytoplasm. TFE3 is found in the nucleus of naïve ESCs, where it sustains the naïve GRN via transcriptional control of *Esrrb* (Betschinger et al. 2013). Induced by a metabolic shift, mTORC1-dependent and mTORC1-independent nutrient-sensing pathways converge to cause TFE3 export from the nucleus, thus contributing to the extinction of the naïve GRN (Betschinger et al. 2013; Villegas et al. 2019). Whether the export of TFE3 and the import of FoxO TFs are coordinated remains an interesting open question.

Once translocated to the nucleus, FoxO TFs play key roles in the establishment of the formative GRN and contribute to the expression of a large number of genes that are themselves required for the exit from naïve pluripotency. Our ChIP-seq experiment revealed that FoxO1 binds to multiple genomic locations even in naïve conditions and that FoxO1 2i targets are part of the core naïve TF-network. Hence, FoxO1 could be necessary for the maintenance of naïve identity by directly regulating the naïve TF-network. However, FoxO TFs remain bound to core naïve genes even 24h after the onset of differentiation, when most of the core naïve genes have been transcriptionally inactivated. These binding dynamics pose the question of whether FoxO TFs can act as activators or repressors depending on the cellular context. This suggestion is consistent with the observed downregulation of naïve genes upon FoxO1 nuclear overexpression, the increased levels of FoxO-TF bound core naïve genes after FoxO1 siRNA treatment, and the increased expression of naïve specific genes upon AKT inhibition and subsequent nuclear translocation of FoxO TFs.

But how can such a silencing function be reconciled with the fact that FoxO TFs are mainly known as activators of gene expression? In a recent study on human transcriptional effector domains it was shown that FoxO1 can also display repressor activity (DelRosso et al. 2023). Whether FoxO TFs act as activators or repressors might depend on cellular environment, post-translational modifications on FoxO TFs, or on the co-binding of other TFs and co-factors. This potential dual role of FoxO TFs in mediating both activation and silencing will be an exciting question for future research.

Our data show that a large number of FoxO TF-bound genomic regions are also bound by ESRRB, OCT4 and OTX2. This places FoxO TFs as a core component of both naïve and formative GRNs. Which other factors are involved in initiating the formative GRN remains unclear. β-catenin or Tcf TFs represent attractive candidates. β-CATENIN was found to physically interact with FOXO4 (Bourgeois et al. 2021; Doumpas et al. 2019), thereby increasing the activity of FoxO (Essers et al. 2005). In contrast, interaction between FoxO and TCF transcription factor derived peptides was reported to disrupt β-CATENIN condensate formation, thereby interfering with β-catenin-driven gene expression (Gui et al. 2023). Whether such interactions also exist in pluripotency transitions, and if they contribute to the FoxO TF-dependent role in establishing the formative GRN, remains unclear. However, the significant overlap of β-CATENIN and FoxO1 peaks in our analysis indicates a functional interaction on chromatin.

It is also tempting to speculate about a function for phosphorylated cytoplasmic FoxO TFs, which are abundant in the naïve state in 2i. Cytoplasmic phosphorylated FoxO TFs can bind to the scaffold protein IQGAP1. This interaction prevents IQGAP1-mediated ERK-activation (Pan et al. 2017). Such an interaction would enable cytoplasmic phospho-FoxO to stabilise naïve identity by inhibiting ERK signalling. Nuclear translocation of FoxO TFs at the onset of differentiation would release ERK inhibition. Hence, the contribution of FoxO shuttling to the transition into formative pluripotency could be a dual one: Firstly, transcriptional control of the naïve and formative core GRNs, and secondly by allowing ERK to exert its crucial function at the exit from naïve pluripotency. This would position FoxO TFs as a central cell fate switch that can both shield naïve identity and disrupt it under differentiation-permissive conditions.

FoxO TFs are reported inhibitors of reprogramming to a naïve pluripotent ESC state (Fu et al. 2021; Yu et al. 2014). Our data indicate that the molecular reason for this effect could lie in the post-implantation-fate inducing function of FoxO TFs.

FoxO TFs are classically seen as tumour suppressors, but a pro-tumorigenic role has been proposed (Hornsveld et al. 2018; Jiramongkol and Lam 2020). Whether the control of cell fate specific programmes in response to a shift in signalling states, as reported here, contributes to the role of FoxO TFs in tumour biology remains to be investigated.

In sum, our work highlights a previously unappreciated role of AKT in maintenance of pluripotency by ensuring cytoplasmic sequestration of FoxO TFs. Upon nuclear translocation, FoxO TFs play a key role in the shutdown of the naïve pluripotent identity and the initiation of the formative gene expression programme.

Our findings support the generalization of a paradigm where relocation of TFs regulates the delicately balanced GRNs that govern stem cell transitions.

## Author contribution

LS designed and performed wet-lab and computational experiments, designed the study and wrote the manuscript. SK designed and performed siRNA experiments. GS and NR performed embryo IF stainings. ML designed and supervised the study, and wrote the manuscript.

## Acknowledgements

We thank Kitti Dóra Csályi and Thomas Sauer at the Max Perutz Laboratories FACS Facility for expert support. Next generation sequencing was performed at the Vienna Biocenter Core Facilities (VBCF), and we thank Thomas Grentzinger (VBCF) for the ATAC-seq sample processing. We thank Niklas Urbanek, Michael Bindl, Mattia Pitasi, Michelle Huth, Luis Miguel Alvan Cerron, Julia Ramesmayer, Ana Maria Sandru and Stefania Bucconi for experimental support. We thank all members of the Leeb and Bücker labs for helpful discussions and Manuela Baccarini and Thomas Leonard for support and helpful suggestions. Laura Santini is an OEAW doc.fellowship awardee (DOC/25622) and a member of the FWF-funded doctoral programme ‘Signalling Molecules in Cellular Homeostasis’ (SMICH; W1261). Martin Leeb is a member and speaker of SMICH and a ‘Wiener Wissenschafts-Forschungs- und Technologiefonds (WWTF) Vienna Research Group Leader (VRG14-006).

## Competing interests

The authors declare no competing interests.

## MATERIALS & METHODS

### Cell culture

Mouse embryonic stem cells (mESCs) were cultured on gelatin-coated (Sigma-Aldrich, G1890) plates in DMEM high-glucose medium (Sigma-Aldrich, D5671) supplemented with 10% FBS (Gibco, 10270-106), 2 mM L-Glutamine (Sigma-Aldrich, G7513), 0.1 mM NEAA (Sigma-Aldrich, M7145), 1 mM Sodium Pyruvate (Sigma-Aldrich, S8636), 10 µg/ml penicillin-streptomycin (Sigma-Aldrich, P4333), 55 µM β-mercaptoethanol (Fisher-Scientific, 21985-023), 10 ng/ml LIF (batch tested, in-house) and 2i (1.5 μM PD0325901 and 3 μM CHIR99021), referred to here as ES DMEM-2i medium. mESCs were passaged every second day and routinely tested negative for mycoplasma infection.

All cell lines used in this study were derived from a parental cell line carrying a Rex1-GFPd2 reporter (destabilised version of the GFP transgene (GFPd2) under control of the endogenous Rex1 promoter, (Wray et al. 2011)) and a Cas9 transgene (EF1alpha-Cas9 cassette targeted to the Rosa26 locus, (Li et al. 2018) (RC9 cells). Cells lacking *Pten*, *Tsc2* or *Tcf7l1* (*Pten* KO #1, *Tsc2* KO and *Tcf7l1* KO) had been previously generated (Lackner et al. 2021). An additional *Pten* KO clone (#2) was generated during this study, following the same procedure described in Lackner et al. 2021. Pten rescue cell lines were generated by cloning the PTEN coding sequence (amplified by PCR from RC9 genomic DNA) into a pCAG-3xFLAG-empty-pgk-hph vector (Betschinger et al. 2013).

### Monolayer differentiation

mESCs were plated on gelatin-coated plates at a final density of 1 × 10^4^ cells/cm^2^ in N2B27 medium (1:1 mix of DMEM/F12 (Gibco, 21331020) and Neurobasal medium (Gibco, 21103049) supplemented with N2 (homemade), B27 Serum-Free Supplement (Gibco, 17504-044), 2 mM L-Glutamine (Sigma-Aldrich, G7513), 0.1 mM NEAA (Sigma-Aldrich, M7145), 10 µg/ml penicillin-streptomycin (Sigma-Aldrich, P4333), 55 µM β-mercaptoethanol (Fisher-Scientific, 21985-023)) and 2i (1 μM PD0325901 and 3 μM CHIR99021), hereby referred to as N2B27-2i medium. The following day, cells were washed with PBS and medium was exchanged to either N2B27 without 2i to induce differentiation for the indicated time, or to fresh N2B27-2i for the undifferentiated controls.

### Rapamycin treatment

mESCs were plated on gelatin-coated plates in N2B27-2i + DMSO (Sigma-Aldrich, D2650) or N2B27-2i + 20 nM Rapamycin (Enzo Life Sciences, BML-A275-0005). The following day, cells were washed with PBS and medium was exchanged to N2B27+DMSO or N2B27+ 20 nM Rapamycin to induce differentiation for the indicated time.

### MK-2206 treatment

mESCs were plated on gelatin-coated plates in N2B27-2i or N2B27-2i + 1 μM MK-2206 (Cayman Chemical, 11593). The following day, cells were washed with PBS and medium was exchanged to either N2B27 or N2B27+1 μM MK-2206 to induce differentiation for the indicated time, or to N2B27-2i or N2B27-2i+1 μM MK-2206 for the undifferentiated controls.

### RNAi assay

mESCs were plated on gelatin-coated plates in N2B27-2i and transfected with FlexiTube siRNAs (Qiagen) using DharmaFECT 1 (Fisher Scientific, T-2001). The next day, cells were washed with PBS and medium was exchanged to either N2B27 without 2i to induce differentiation, or fresh N2B27-2i for the undifferentiated controls. Two siRNAs targeting FoxO1 (SI01005200, SI02694153) were used at a final concentration of 20 nM. As controls, siRNAs targeting GFP (siGFP) or scrambled siRNAs (siScr) were used.

### Expression of nuclear FoxO1

A FoxO1 coding sequence carrying three mutations (T24A, S253D and S316A) was cloned from the an addgene vector (Plasmid #12149) (Kitamura et al. 2005) into a pB-TetOn-3xFLAG-Empty-PolyA-Puro vector. The resulting plasmid, hereby referred to as 3xFLAG-FoxO1^nuc^, was transfected into RC9 and *Pten* KO cells. Single-cell derived, independent clones were selected and expanded for further experiments. For experiments in differentiation-permissive conditions, RC9-based and *Pten* KO-based clones were plated on gelatin-coated plates in N2B27-2i. The following day, cells were washed with PBS and medium was exchanged to N2B27 with or without 500 ng/ml of Doxycycline (Sigma-Aldrich, D9891). After 8 hours, medium was changed to N2B27 and cells were differentiated for further 16 hours. For the experiments in naïve conditions, RC9-based clones were plated in N2B27-2i and cultured for 48 hours. The last 8 hours before harvesting, 500 ng/ml of Doxycycline was added to the medium.

### Flow Cytometry analysis

After the indicated amount of time in differentiation (N2B27-based) or control (N2B27-2i-based) medium conditions, cells were harvested using 0.25% trypsin/EDTA and resuspended in ES-DMEM medium to neutralise trypsin. Rex1-GFPd2 levels were measured using the LSRFortessa flow cytometer (BD bioscience) and then analysed with the FlowJo software (v10, BD bioscience).

### Real-time PCR analysis

After the indicated amount of time in differentiation (N2B27-based) or control (N2B27-2i-based) medium conditions, cells were washed with PBS and harvested in RNA Lysis buffer containing 1% (v/v) 2-mercaptoethanol and stored at −80°C before isolation of RNA. RNA was extracted using the ExtractMe kit (Blirt, EM15) following the manufacturer’s instructions. 0.25– 1 µg of RNA was reverse-transcribed into cDNA using the SensiFAST cDNA Synthesis Kit (Bioline, BIO-65054). Real-time PCR was performed on the CFX384 Touch real-time PCR detection system (Bio-Rad) using the Sensifast SYBR no Rox kit (Bioline, BIO-98020). Data analysis and visualisation was performed using Microsoft Excel (Office 365) and R. Used primers are listed in *Supplementary Table 3*.

### Immunoblotting analysis

After the indicated amount of time in differentiation (N2B27-based) or control (N2B27-2i-based) medium conditions, cells were washed with PBS and harvested in RIPA buffer (Sigma-Aldrich, 20-188) supplemented with Complete Mini EDTA-free protease inhibitor cocktail (Roche, 04693159001) and PhosSTOP (phosphatase inhibitor cocktail (Roche, 04906845001). Protein extraction was performed by incubating the samples on ice for 15 minutes and then collecting the supernatant after centrifugation at 13,000 rpm at 4°C for 30 minutes. Protein concentration was determined using a Bradford Assay (Bio-Rad). 8-20 µg whole cell lysates were separated on 8-12% SDS–PAGE gels (depending on the molecular weight of the target proteins) and subsequently blotted on 0.2 µm nitrocellulose membranes (Amersham). Membranes were blocked at RT for 1 hour with 5 % milk diluted in PBS (Sigma-Aldrich, P4417) containing 0.1% Tween-20 (PBS-T). Primary antibodies were incubated overnight at 4°C or for 1 h at room temperature (RT). Secondary antibodies were incubated for 1 h at RT. The following primary antibodies were diluted in PBS-T containing 5% BSA and used 1:1000 for anti-phospho-Akt(Ser473) (rabbit; Cell Signaling, 4058), 1:1000 for anti-phospho-Akt(Thr308) (rabbit; Cell Signaling, 13038), 1:1000 for anti-pan-Akt (rabbit; Cell Signaling, 4691), 1:1000 for anti-phospho-GSK3β(Ser9) (rabbit; Cell Signaling, 9336), 1:1000 for anti-GSK3β (rabbit; Cell Signaling, 12456), 1:1000 for anti-phospho-4E-BP1(Ser65) (rabbit; Cell Signaling, 9451), 1:1000 for anti-4E-BP1 (rabbit; Cell Signaling, 9644), 1:1000 for anti-phospho-p70 S6 kinase(Thr389) (rabbit; Cell Signaling, 9234), 1:1000 for anti-p70 S6 kinase (rabbit; Cell Signaling, 2708), 1:1000 for anti-PTEN (rabbit; Cell Signaling, 9559), 1:1000 for anti-TSC2 (rabbit; Cell Signaling, 4308), 1:1000 for anti-FoxO1 (rabbit; Cell Signaling, 2880) and 1:1000 for anti-FoxO3a (rabbit; Cell Signaling, 2497). The following primary antibodies were diluted in PBS-T containing 5% milk and used 1:1000 for anti-NANOG (rabbit; NovusBio, NB100-58842), 1:5000 for anti-Tubulin (mouse; Sigma-Aldrich, T8203), 1:10,000 for anti-GAPDH(G-9) (mouse; Santa Cruz Biotechnology, sc-365062) and 1:10,000 for anti-Vinculin(H-10) (mouse; Santa Cruz Biotechnology, sc-25336). Secondary antibodies were diluted in 5% milk and used 1:10,000 for anti-rabbit IgG (Amersham, NA934) and 1:15,000 for anti-mouse IgG (goat; Santa Cruz Biotechnology, sc-2064). Chemiluminescence signal was detected using the ECL Select detection kit (GE Healthcare, GERPN2235) with a ChemiDoc system (Bio-Rad). Data analysis was performed using ImageLab.

### Immunofluorescence analysis

mESCs were plated at a final density of 1 × 10^4^ cells/cm^2^ on fibronectin-coated (MM Merck Millipore, FC010) µ-Slide 8 Well Glass Bottom Chamber Slides (Ibidi, 80827) in N2B27-2i medium. The following day, cells were washed with PBS and medium was exchanged to either N2B27 without 2i to induce differentiation, or to fresh N2B27-2i for the undifferentiated controls. After the indicated amount of time, cells were washed with PBS and fixed for 15 minutes at RT with freshly diluted 4% PFA (16% paraformaldehyde diluted in 1:4 in PBS) (SCI Science Services, E15710). Cells were washed in PBS and subsequently permeabilized with PBS containing 0.1% Triton-X for 10 minutes at RT. Cells were washed 3x in PBS-T and then blocked using PBS-T containing 5 % BSA (blocking buffer) for 30 minutes at 4°C. Primary antibodies were incubated overnight at 4°C. Cells were washed 3x with PBS-T. Secondary antibodies were incubated for 1 hour at RT. Cells were washed 2x with PBS-T. Nuclei were stained with 1 µg/ml DAPI (Sigma-Aldrich, D9542) for 10 minutes at RT. Cells were washed 3x with PBS and stored in PBS at 4°C until the image acquisition procedure. Primary and secondary antibodies were diluted in blocking buffer and used 1:100 for anti-FoxO1 (rabbit; Cell Signaling, 2880), 1:100 for anti-ESRRB (mouse; R&D Systems, PP-H6705-00), 1:250 for anti-FLAG M2 (mouse; Sigma-Aldrich F1804), 1:200 for anti-NANOG (rabbit; NovusBio, NB100-58842), 1:500 for anti-mouse Alexa-555 (donkey; Cell Signaling, 4409), 1:500 for anti-rabbit Alexa-647 (goat; Cell Signaling, 4414). Images were acquired using a Zeiss LSM 980 confocal microscope. Images were analyzed using Fiji/ImageJ. For quantifying the nuclear intensity of the stained proteins, segmentation was performed on the DAPI channel with cellpose with the following settings: cell diameter (in pixels) = 170, flow_threshold = 0.4, cellprob_threshold = 0.0, stitch_threshold = 0.0, model = cyto2. The obtained nuclei outlines were imported into Fiji/ImageJ and used to create mask objects. The latter were used to measure the mean fluorescence intensity of each nucleus in all recorded channels (besides the DAPI channel). Nuclei out of focus were manually removed from the analysis. Data analysis and visualisation was subsequently performed in R.

### Intracellular staining

After the indicated amount of time in differentiation (N2B27-based) or control (N2B27-2i-based) medium conditions, cells were harvested using 0.25% trypsin/EDTA and resuspended in ES-DMEM medium to neutralise trypsin. Cells were centrifuged, and cell pellets were washed twice with PBS before fixation with 2% PFA for 15 minutes at RT. Cells were washed with FACS buffer (PBS containing 5% BSA) and subsequently permeabilized with ice-cold MeOH for 10 minutes on ice. After 3 washes in FACS buffer, cells were incubated in FACS buffer for 10 minutes in the dark. Subsequently, cells were incubated with primary antibodies for 1 hour at RT. Cells were then washed 3x with FACS buffer, and incubated with secondary antibodies for 15 minutes on ice. Cells were washed 3x with FACS buffer, and stored in FACS buffer until flow cytometry analysis at the LSRFortessa flow cytometer. Primary and secondary antibodies were diluted in FACS buffer and used 1:100 for anti-phospho-Akt(Ser473) (rabbit; Cell Signaling, 4058) and 1:500 for anti-rabbit Alexa-647 (goat; Cell Signaling, 4414). Flow cytometry data was analysed with the FlowJo software. Mean fluorescence intensities (MFI) for stained samples were calculated by subtracting the MFI of their relative controls (cells stained only with the secondary antibody). Data analysis and visualisation was performed in R.

### Nucleo-cytoplasmic fractionation

Subcellular fractionation experiments were performed following a protocol adapted from Rockland (https://www.rockland.com/resources/nuclear-and-cytoplasmatic-extract-protocol/). 1.0 × 10^4^ cells/cm^2^ of WT and *Pten* KO cells were plated in 10 cm gelatine-coated plates. The next day, cells were washed with PBS and medium was exchanged to either N2B27 without 2i to induce differentiation, or to fresh N2B27-2i for the undifferentiated controls. After 24 hours cells were harvested. 1/20 of cells were processed as described above for total protein extraction. The rest was used for the nucleo-cytoplasmic fractionation. Cell pellets were resuspended in 5 pellet volumes of CE buffer adjusted to pH 7.6 (10 mM HEPES, 60 mM KCl, 1 mM EDTA, 0.075% NP-40, 1mM DTT, 1 mM PMSF) and incubated on ice for 3 minutes. Samples were pelleted by centrifugation at 1300 rpm for 4 minutes at 4°C and the supernatant (the cytoplasmic fraction) was transferred to clean tubes. The nuclei were washed carefully with 5 pellet volumes of CE buffer (without NP-40), and then pelleted at 1300 rpm for 4 minutes at 4°C. The supernatant was discarded, and the nuclei were resuspended in 1 pellet volume of NE buffer adjusted to pH 8.0 (20 mM Tris Cl, 420 mM NaCl, 1.5 mM MgCl2, 0.2 mM EDTA, 1mM PMSF, 25% glycerol) and the salt concentration was then adjusted to 400 mM with 5 M NaCl. An additional pellet volume of NE buffer was added to the extracts before incubating them for 10 minutes on ice. The extracts were vortexed every 3 minutes during the incubation time. Both cytoplasmic and nuclear fractions were spun at maximum speed to pellet any remaining nuclei. Cytoplasmic and nuclear fractions were transferred to clean tubes, glycerol was added to the cytoplasmic fraction to 20%. Both fractions were stored at −80°C.

### RNA-sequencing analysis

For differentiation experiments using RC9, *Pten* KO and *Tsc2* KO cells, and for the 2h-resolved time course, count tables generated in a previous study were used (Lackner et al. 2021). QuantSeq analysis was performed for all the other RNA-seq experiments described in the manuscript. For the Rapamycin experiment, WT, *Pten* KO and *Tsc2* KO cells treated with DMSO or Rapamycin from two independent differentiation assays were sequenced (duplicates). For the FoxO1 knockdown experiment, WT cells were treated either with an individual siRNA sequence targeting *Foxo1* transcript, or with a combination of both siRNAs (triplicates, siRNA #1, siRNA#2, siRNA#1 + siRNA#2). For the control sample (siCtrl), two siGFP samples combined with 1 siScr sample were considered a triplicate. For the MK-2206 experiment, WT cells left untreated or treated with MK-2206 from two independent differentiation assays were sequenced (duplicates). Library preparation (according to the Lexogen 3’ mRNA Seq Library Prep Kit), multiplexing (by qPCRs) and sequencing on an Illumina NextSeq2000 P3 platform was carried out at the VBC NGS facility. 5-10 million of single end reads at 50 bp read length were generated per sample. The resulting fastq files were analysed with a Nextflow 23.04.1.5866 /nf-core/rnaseq v3.10.1 pipeline. Quality control was performed using fastQC (v0.11.9), and transcripts were mapped to the mm10 assembly mouse reference genome using Salmon (v1.9.0) as a pseudoaligner and STAR (v2.7.10a) as an aligner. DESeq2 (v1.38.3) was used to generate normalised count tables and to perform differential expression analyses (FDR-adjusted *p-value* ≤ 0.05; H0: log2FC = 0). pheatmap (v1.0.12), EnhancedVolcano (v1.16.0), UpsetR (v1.4.0), eulerr (v7.0.0) and ggplot2 (v3.4.0) were used for data visualisation in R. Combined lists of upregulated and downregulated genes in *Pten* KO and *Tsc2* KO were generated by selecting genes differentially expressed (DEGs) in both KOs (log2FC ≥ 0.5 for the upregulated genes, and log2FC ≤ −0.5 for the downregulated genes).

### Chromatin Immunoprecipitation (ChIP)

FoxO1 and FoxO3 ChIP were performed as described in Thomas et al. 2021. 1.5 × 10^4^ cells/cm^2^ of WT and *Pten* KO cells were plated in duplicate on 15 cm gelatine-coated plates. The next day, cells were washed with PBS and medium was exchanged to either N2B27 without 2i to induce differentiation, or to fresh N2B27-2i for the undifferentiated controls. After 24 hours, cells were harvested. Cells were washed with PBS, and then cross-linked directly on the plate with 1% formaldehyde in PBS for 10 minutes. Subsequently, 0.125 M glycine was added to the plates for 10 minutes to quench cross-linking. The plates were washed 2x with PBS, and then cells were scraped off in ice-cold PBS containing 0.01% Triton-X. Cells were pelleted by centrifugation at 500 g for 5 minutes and flash-frozen in liquid nitrogen. Cell pellets were resuspended in 5 ml LB1 (50 mM Hepes pH 7.5, 140 mM NaCl, 1 mM EDTA, 10% glycerol, 0.5% NP-40, 0.25% TX-100, 1 mM PMSF, 1×Complete Mini EDTA-free protease inhibitor cocktail) to extract nuclei and rotated vertically for 10 minutes at 4°C. Nuclei were pelleted by centrifugation at 1350 g for 5 minutes at 4°C, and then resuspended in 5 ml LB2 (10 mM Tris pH 8.0, 200 mM NaCl, 1 mM EDTA, 0.5 mM EGTA1, mM PMSF, 1xComplete Mini EDTA-free protease inhibitor cocktail), and rotated vertically for 10 minutes at RT. Samples were pelleted by centrifugation at 1350 g for 5 minutes at 4°C and then resuspended in 1.5 ml LB3 (10 mM Tris–HCl pH 8.0, 100 mM NaCl, 1 mM EDTA 0.5 mM EGTA, 0.1% Na-deoxycholate, 0.5% N-lauroylsarcosine, 1 mM PMSF, 1xComplete Mini EDTA-free protease inhibitor cocktail) and 200 μl sonification beads (diagenode) in Bioruptor® Pico Tubes (diagenode). Chromatin was sonicated for 13 cycles with 30 seconds on, 45 seconds off parameters. Sonicated samples were transferred to fresh tubes and centrifuged at 16’000 g at 4°C to pellet cellular debris. 1.1 ml of supernatant were collected and transferred to fresh tubes. 110 μl of 10% Triton-X were added to a final concentration of 1%. For each sample, 50 μl were collected as input and 1 ml was used for immunoprecipitation. Chromatin was incubated with 20 μl (1:50 dilution) of anti-FoxO1 (rabbit; Cell Signaling, 2880) or anti-FoxO3a (rabbit; Cell Signaling, 2497) antibodies overnight at 4°C with vertical rotation. Samples were collected after 14 hours. 100 μl of Dynabeads protein G (Thermo Fisher Scientific, 10765583) per sample were washed in ice-cold blocking solution (PBS containing 0.5% BSA) and then incubated with the antibody-bound chromatin solutions for 4 hours. Beads were washed 5x in ice-cold RIPA wash buffer (50 mM Hepes pH 7.5, 500 mM LiCl, 1 mM EDTA, 1% NP-40, 0.7% Na-Deoxycholate), and then 3x with TE + 50 mM NaCl. Samples were eluted in 210 μl elution buffer (50 mM Tris pH 8.0, 10 mM EDTA, 1% SDS) for 15 min at 65°C. Supernatant containing the antibody-bound chromatin fraction was separated from the beads. Three volumes of elution buffer were added to the Input samples. Decrosslinking was performed by incubating both Input and ChIP samples at 65°C overnight. The next day, one volume of TE and RNase A (to a 0.2 mg/ml final concentration) were added to the Input and ChIP samples, followed by incubation for 2 hours at 37°C. Final salt concentration was adjusted to 5.25 mM CaCl_2_ with 300 mM CaCl_2_ in 10 mM Tris pH 8.0. Samples were then incubated with 0.2 mg/ml Proteinase K for 30 minutes at 55°C. DNA was extracted with phenol-chloroform using Phase Lock GelTM tubes (Quantabio, 733-2478) and then precipitated in EtOH. DNA pellets were dissolved in H_2_O.

### ChIP-sequencing analysis

Libraries were prepared at the VBC NGS facility and sequenced on an Illumina NovaSeq platform. Within the FoxO1 ChIP experiment, 20-40 million paired end reads at 150 bp read length were generated for the Input samples, and 80-130 million reads for the ChIP samples. Within the FoxO3 ChIP experiment, ∼ 25 million single end reads at 100 bp read length were generated for both Input and ChIP samples. RC9-based FoxO3 ChIP samples were re-sequenced to increase sequencing depth, and an additional ∼ 60 million of paired end reads at 150 bp read length per sample were generated. R1 reads from the two sequencing runs were concatenated before data processing. FoxO1 ChIP and Input reads were processed following a paired end mode, while FoxO3 ChIP and Input samples following a single end mode. Quality control of fastq files was performed using fastQC (v0.11.9) before and after trimming the sequencing adapter fragments with trim-galore (v0.6.7) and cutadapt (v3.5). Additional 2 bp at the 3’ end were removed with the parameters *--three_prime_clip_R1 2* (*--three_prime_clip_R2 2* in case of paired end reads). The trimmed reads were aligned to the mm10 assembly mouse reference genome with bowtie2 (v2.4.4), with an alignment rate of 80-98%. The obtained sam files were converted to bam files with samtools (v1.13), and uniquely mapping reads were extracted by removing duplicate reads (as potential PCR artefacts) with samtools *markdup*. Peak calling was performed on bam files with macs2 (v2.2.7.1) on combined ChIP duplicates, using all Input samples as control files. Potential artefactual regions listed in the mm10 blacklist (http://mitra.stanford.edu/kundaje/akundaje/release/blacklists/mm10-mouse/mm10.blacklist.bed.gz) were removed from the obtained bed files using bedtools (v2.31.0). Peaks assigned to unidentified regions (chrUn) were manually removed.

### ATAC-sequencing analysis

1.0 × 10^4^ cells/cm^2^ cells were plated on gelatine-coated plates. The next day, cells were washed with PBS and medium was exchanged to either N2B27 without 2i to induce differentiation, or to fresh N2B27-2i for the undifferentiated controls. After 24 hours cells were harvested and counted. 250,000 cells per sample were submitted to the VBC NGS facility for further processing and ATAC-seq library preparation (Bulk ATAC-seq Illumina). In brief, cells were lysed with 0.5x lysis buffer (0.01 M Tris-HCl pH 7.5, 0.01 M NaCl, 0.003M MgCl_2_, 1 % BSA, 0.1% Tween-20, 0.05% NP-40, 0.005% Digitonin, 0.001 M DTT, 1 U/μl RNAse inhibitor), and tagmentation reaction (Tn5 Illumina) was performed on 50,000 isolated nuclei. Libraries were prepared with the Nextera DNA Library Preparation kit and sequenced on an Illumina NovaSeq platform. 80-170 million of paired end reads at 150 bp read length were generated per sample. The resulting fastq files were analysed with a Nextflow 23.04.1.5866 /nf-core/atacseq v2.0 pipeline. Quality control was performed using fastQC (v0.11.9), and reads were mapped to the mm10 assembly mouse reference genome using bowtie2 (v2.4.4). Peak calling was performed on the bam files generated by the nfcore pipeline with Genrich (v0.6.1). Consensus peaks were defined as peaks with at least a 50% overlap between replicates and generated with samtools *intersect*. Bed files were filtered for the mm10 blacklist using bedtools. Peaks assigned to unidentified regions (chrUn) were manually removed.

### Motif enrichment analyses

Motif enrichment analysis was performed with Homer (v4.11). For finding motifs enriched in FoxO1 or FoxO3-bound regions, *findMotifsGenome.pl* was used with default settings.

For finding enriched motifs in ESCs or EpiLCs enhancers, enhancer lists from the Bücker lab were used (filtered to exclude TSS, Thomas et al. 2021). A list of overlapping enhancers between ESCs and EpiLCs was generated with bedtools and used as background to identify ESC-specific and EpiLC-specific enriched motifs with *findMotifsGenome.pl* using default settings. According to Homer’s documentation, a motif can be considered enriched when its associated *p-value* is lower than 1e-50.

### Data integration analyses

Downstream analyses for ChIP-seq and ATAC-seq were performed with Deeptools (v3.5.1). BigWigs were generated from single bam files with *bamCoverage* using a binsize of 10 bp and a normalisation coverage to 1x mouse genome size (RPGC) excluding the X chromosome. Principal component analysis (PCA) was performed with *multiBigwigSummary* and *plotPCA*. Bigwigs replicates were merged using *bigWigMerge*, and heatmaps were plotted on merged BigWigs using *computeMatrix* and *plotHeatmap*. Heatmaps were sorted based on FoxO1 or FoxO3 signals. 3 lists of peaks (separately for FoxO1 and FoxO3 ChIP) were generated with bedtools by intersecting WT (RC9) bed files: 2i-only peaks, N24-only peaks and shared peaks. Peak to gene association was performed on these peak lists in R with ChIPSeeker (v1.34.1). Oct4 and Otx2 ChIP BigWigs and peak lists were obtained from the Bücker lab (Buecker et al. 2014). Esrrb ChIP BigWigs and bed files were obtained from the Martello lab (Carbognin et al. 2023). β-catenin ChIP bed files were downloaded from CODEX (https://codex.stemcells.cam.ac.uk/), and mm9 coordinates were converted into mm10 coordinates using the liftOver tool from UCSC (https://genome.ucsc.edu/cgi-bin/hgLiftOver).

Overlap between genomic regions was performed with ChIPPeakAnno (v3.32.0). For the FoxO1-FoxO3 ChIP overlap, the background for the hypergeometric test was set to the total number of detected open chromatin regions (generated by merging ATAC-seq peaks in 2i with ATAC-seq peaks at N24 with ESC). For all other overlaps, the background used was obtained by merging all ATAC-seq peaks with the lists of ESC and EpiLC enhancers (Thomas et al. 2021). Genome tracks shown in Figure 4 were generated using karyoploteR (v1.24.0) in R.

### Enrichment analyses

All enrichment analyses presented in this work were performed in R with hypeR (v2.0.1). For KEGG pathway enrichment analysis, mouse KEGG database was downloaded from http://rest.kegg.jp/link/mmu/pathway and transformed into a hypeR-compatible gene set using the gsets function. As background, the list of all detected genes in the relative RNA-seq experiment was used.

Custom gene sets were also generated with the gsets function. As background for FoxO TF-based enrichments, a list of genes associated with all open chromatin regions was used. Enrichment results were visualised with ggplot2.

### Statistical analysis

All statistical analyses were performed in R with the ggpubr (v0.6.0) package. Information on statistical tests and replicate numbers are provided in the figure legends. Wherever necessary, correction for multiple testing was performed.

---

**Supplementary Fig. 1.**
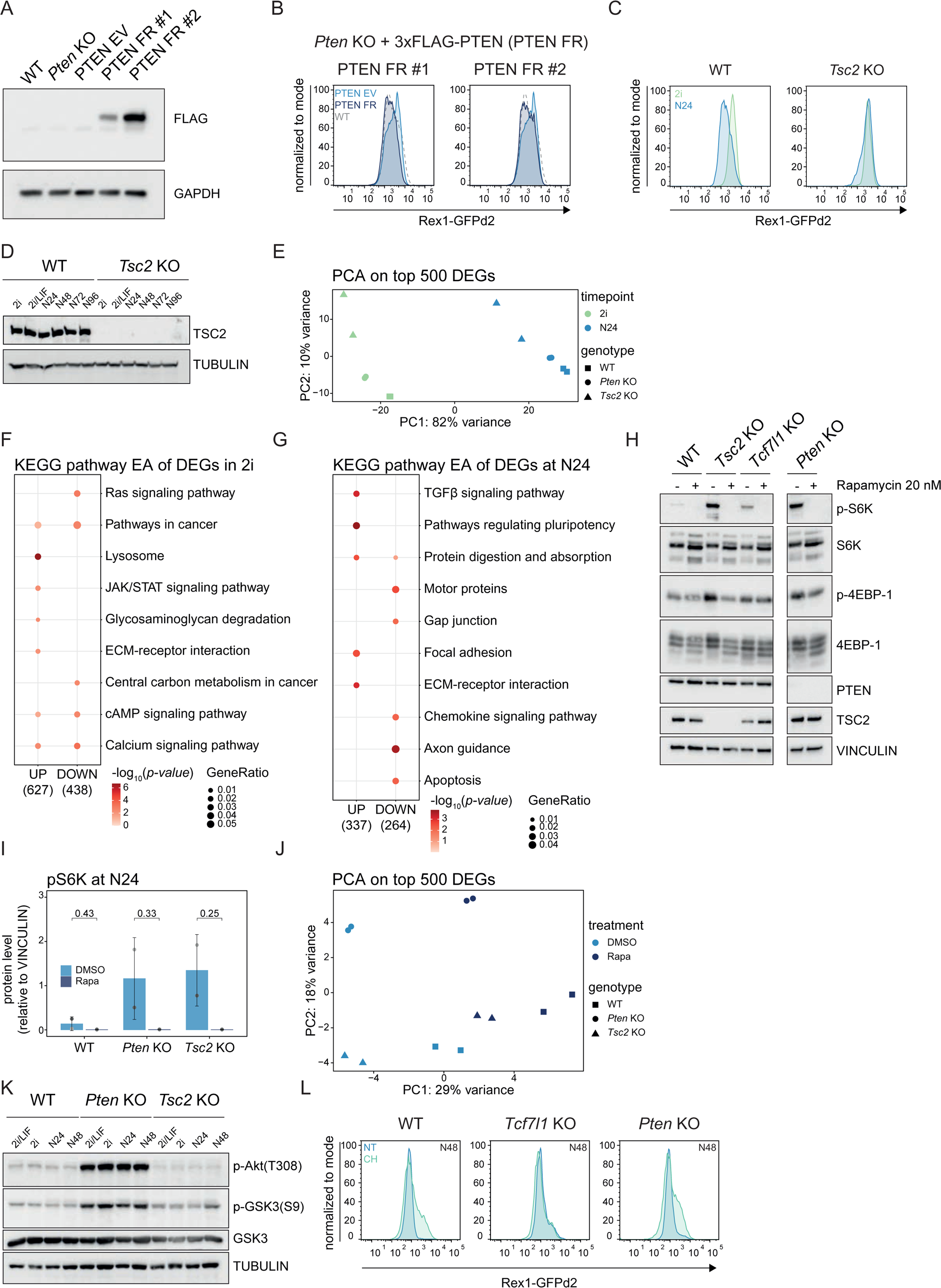
mTORC1 inhibition only partially rescues the differentiation defect of *Pten* KO ESCs. A, Western blot analysis for FLAG expression in WT, *Pten* KO, and PTEN rescue mESCs (FLAG-rescue (FR) or empty vector control, PTEN EV). TUBULIN was used as a loading control. B, Flow cytometry analysis of Rex1-GFP levels in WT (grey dashed line), PTEN EV (light blue profiles) and PTEN FR mESCs (dark blue profiles) at N24. One representative of n=3 independent experiments is shown. C, Flow cytometry analysis of Rex1-GFP levels in WT and in *Tsc2* KO cells in 2i (green profiles) and at N24 (light blue profiles). One representative of n >10 independent experiments is shown. D, Western blot analysis for TSC2 expression in WT and *Tsc2* KO cells in naïve pluripotency supporting conditions (2i and 2i/LIF) and 24h (N24), 48h (N48), 72h (N72) and 96h (N96) after 2i removal. TUBULIN was used as a loading control. E, Principal component analysis (PCA) based on the top 500 DEGs in RNA-Seq data of WT, *Pten* KO and *Tsc2* KO mESCs in 2i (green) and at N24 (light blue). F, KEGG pathway enrichment analysis (EA) on combined lists of upregulated (UP) and downregulated (DOWN) genes in *Pten* and *Tsc2* KO mESCs (x axis). Top 5 categories enriched in each list are shown on the y axis. Dot colour indicates *p-values* (only *p* ≤ 0.1 are shown). Dot size indicates the GeneRatio (ratio between the overlap size and the category size). G, Similar to F for DEGs at N24. H, Western blot analysis for the indicated proteins in WT, *Pten* KO, *Tsc2* KO and *Tcf7l1* KO mESCs at N24 after treatment with DMSO (−) or 20 nM Rapamycin (+). VINCULIN was used as a loading control. One representative of n=2 independent experiments is shown. I, Quantification of pS6K protein expression measured by Western blot analysis in WT, *Pten* KO and *Tsc2* KO cells at N24 after treatment with DMSO (light blue) or 20 nM Rapamycin (Rapa, dark blue). Expression was normalised to VINCULIN. Mean and SD for n=2 independent experiments (distinguished by a greyscale) are shown. Indicated *p-values* show results of paired, two-tailed t-tests. J, PCA analysis based on the top 500 DEGs in RNA-Seq data of WT, *Pten* KO and *Tsc2* KO mESCs at N24 after treatment with DMSO (light blue) or 20 nM Rapamycin (Rapa, dark blue). Each symbol refers to one specific cell line as indicated in the legend. K, Western blot analysis for the indicated proteins in WT, *Pten* KO and *Tsc2* KO mESCs in naïve pluripotency supporting conditions (2i and 2i/LIF) and 24h (N24) and 48h (N48) after 2i removal. TUBULIN was used as a loading control. L, Flow cytometry analysis of Rex1-GFP levels in WT, *Tcf7l1* KO and *Pten* KO cells at N48 after treatment with 3 μM CHIRON (CH, green profiles) or left untreated (NT, light blue profiles). One representative of n=2 independent experiments is shown.

**Supplementary Fig. 2.**
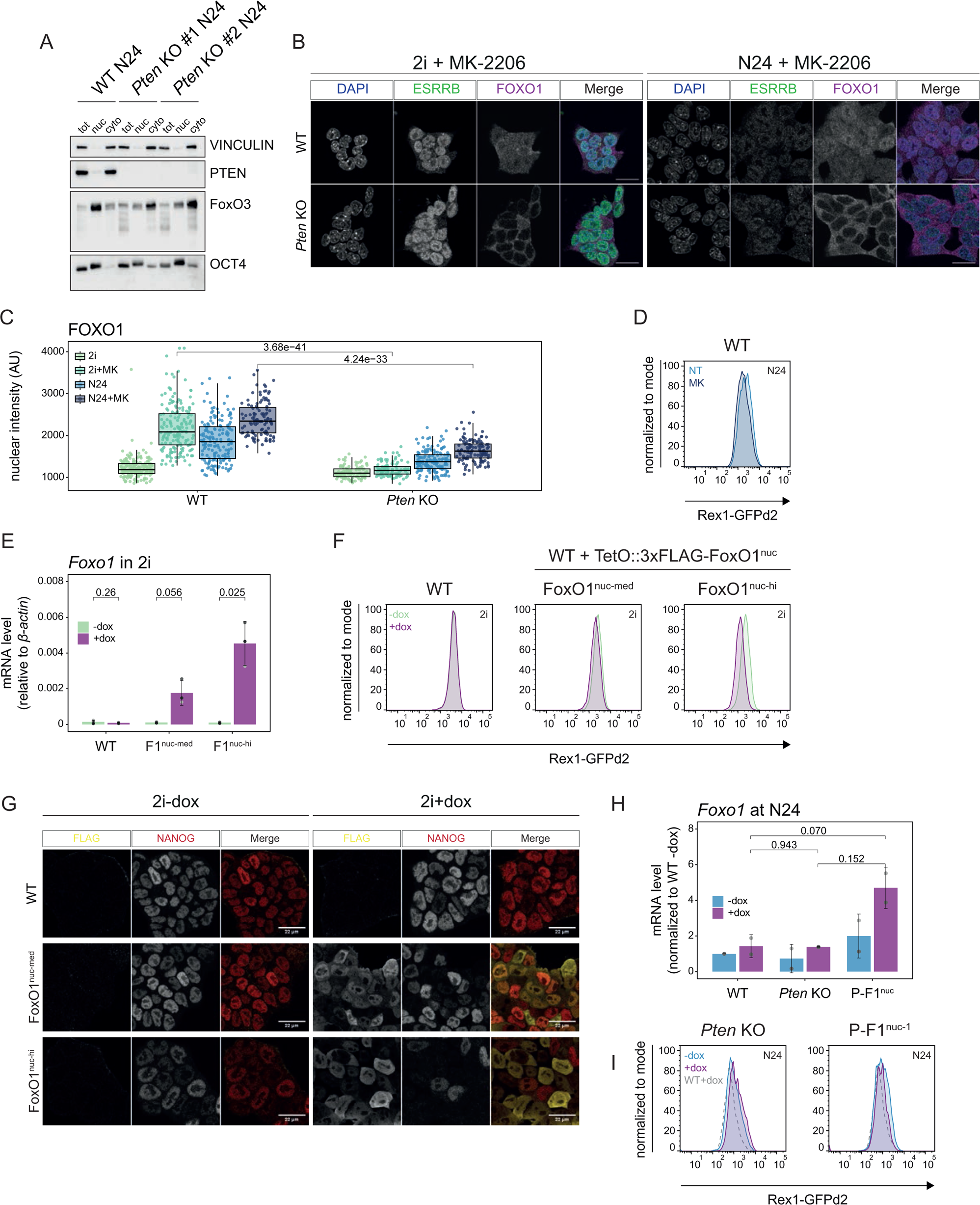
FoxO TFs translocate into the nucleus upon exit from the naive pluripotent state. A, Subcellular localization of the indicated proteins measured by Western blot analysis of nucleo-cytoplasmic fractionation experiments in indicated cell lines at N24. Equal amounts of total (tot), nuclear (nuc) and cytoplasmic (cyt) fractions were loaded. VINCULIN and OCT4 were used as controls for the purity of the cytoplasmic and nuclear fractions, respectively. B, Confocal microscopy images after IF showing FOXO1 (purple) and ESRRB (green) in WT and *Pten* KOs in 2i and at N24 after treatment with 1 μM MK-2206. DAPI staining is shown in blue. One representative of n=2 independent experiments is shown. Scale bar = 20 μM. C, Quantification of FOXO1 nuclear intensity measured from confocal images of untreated WT (n=189 in 2i and n=164 at N24) and *Pten* KO cells (n=100 in 2i and n=153 at N24) as in Fig. 2A and upon treatment with the MK-2206 inhibitor (WT: n=176 in 2i and n=111 at N24; *Pten* KO: n=100 in 2i and n=121 at N24) as in Supplementary Fig. 2B. Data from n=2 independent experiments is shown. The indicated *p-values* show the results of two-tailed Wilcoxon rank sum tests. D, Flow cytometry analysis of Rex1-GFP levels in WT cells 24h after treatment with 1 μM MK-2206 (MK, dark blue profiles) and untreated control (NT, light blue profiles). One representative of n=3 independent experiments is shown. E, Expression levels of *Foxo1* measured by RT-qPCR in WT cells expressing 3xFLAG-FoxO1^nuc^ (FoxO1^nuc-med^ and FoxO1^nuc-hi^) in 2i after 8 hours treatment with 500 ng/ml doxycycline (+dox, purple) or left untreated (-dox, green). Control WT cells are also included. Mean and SD of n=3 independent experiments (distinguished by distinct shades of grey) are shown. Expression was normalised to *β-actin*. *p-values* show results of paired, two-tailed t-tests. F, Flow cytometry analysis of Rex1-GFP levels in WT, FoxO1^nuc-med^ and FoxO1^nuc-hi^ cells in 2i after 8 hours treatment with 500 ng/ml doxycycline (+dox, purple) or left untreated (-dox, green). One representative of n=3 independent experiments is shown. G, Confocal analysis after IF of 3xFLAG-FoxO1 (yellow) and NANOG (red) in WT, FoxO1^nuc-med^ and FoxO1^nuc-hi^ cells in 2i after 8 hours treatment with 500 ng/ml doxycycline (+dox) or left untreated (-dox). H, Expression levels of *Foxo1* measured by RT-qPCR in *Pten* KO cells expressing 3xFLAG-FoxO1^nuc^ (P-FoxO1^nuc^) at N24 after 8 hours treatment with 500 ng/ml doxycycline (+dox, purple) or left untreated (-dox, light blue). Control WT cells and *Pten* KO cells are also included. Mean and SD of n=2 independent experiments (distinguished by a greyscale) are shown. Expression was normalised to *β-actin* and shown as relative to WT in -dox condition. *p-values* show results of paired, two-tailed t-tests. I, Flow cytometry analysis of Rex1-GFP levels in *Pten* KO cells expressing 3xFLAG-FoxO1^nuc^ (P-FoxO1^nuc^) at N24 after 8 hours treatment with 500 ng/ml doxycycline (+dox, purple profiles) or left untreated (-dox, light blue profiles). Rex1-GFP levels of dox-treated WT cells are shown as a grey dashed line. One representative of n=2 independent experiments is shown.

**Supplementary Fig. 3.**
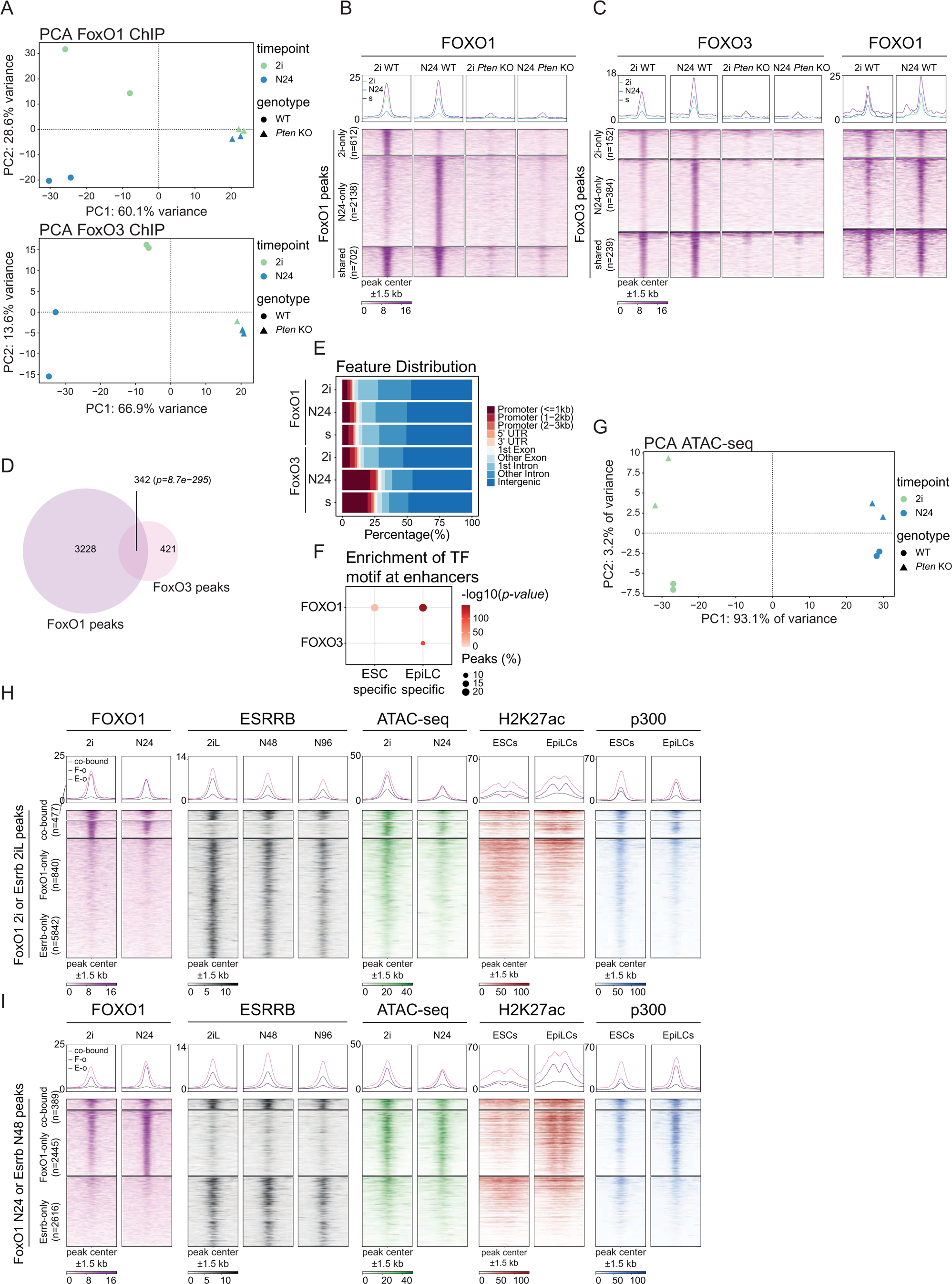
Chromatin dynamics of FoxO TFs at the exit from naïve pluripotency. A, PCA analysis of FOXO1 (top) or FOXO3 (bottom) ChIP-Seq data in WT and *Pten* KO cells in 2i (green) and at N24 (light blue). Each symbol refers to one specific cell line as indicated in the legend. B, Heatmap showing FOXO1 signal in a 1.5 kb window around the center of FoxO1 peaks, divided in 2i-only peaks (n=612, green), N24-only peaks (n=2138, light blue) and shared peaks (n=702, purple), in WT and *Pten* KO cells in 2i or at N24. C, Heatmap showing FOXO3 signal (left) and FOXO1 signal (right) in a 1.5 kb window around the center of FoxO3 peaks, divided in 2i-only peaks (n=152, green), N24-only peaks (n=384, light blue) and shared peaks (n=239, purple), in WT and *Pten* KO cells in 2i or at N24. D, Venn diagram showing the overlap between FoxO1 (purple) and FoxO3 (pink) peaks. The resulting *p-value* from the hypergeometric test of the overlap is shown in italics. E, Feature distribution of FoxO1 and FoxO3 peaks. The categories are indicated in the legend. F, Enrichment analysis of FOXO1 and FOXO3 motifs in ESC- or EpiLC-specific enhancers. Dot colour and size indicate the *p-values* and the percentage of enhancers (“peaks”) with the motif, respectively. G, PCA analysis of ATAC-Seq data in WT and *Pten* KO cells in 2i (green) and at N24 (light blue). Each symbol refers to one specific cell line as indicated in the legend. H, Heatmaps showing FOXO1, ESRRB, ATAC-seq, H3K27ac and p300 signal in the indicated samples in a 1.5 kb window around the center of FoxO1 2i and Esrrb 2iL peaks, divided in co-bound peaks (n=477, pink), FoxO1-only peaks (n=840, purple) and Esrrb-only peaks (n=5842, grey). I, Heatmaps showing FOXO1, ESRRB, ATAC-seq, H3K27ac and p300 signal in the indicated samples in a 1.5 kb window around the center of FoxO1 N24 and Esrrb N48 peaks, divided in co-bound peaks (n=389, pink), FoxO1-only peaks (n=2445, purple) and Esrrb-only peaks (n=2616, grey).

**Supplementary Fig. 4.**
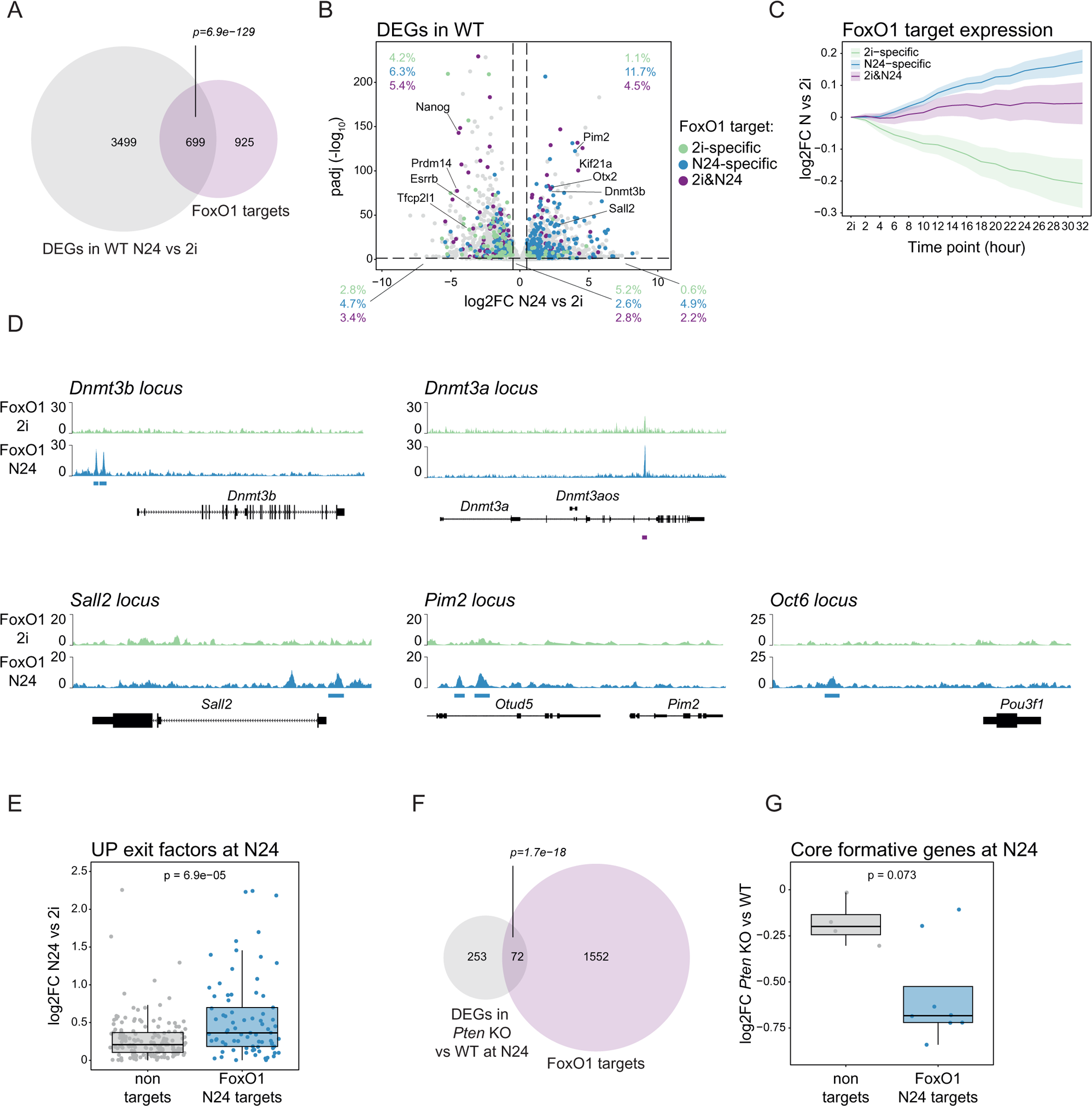
FoxO TF targets are key players in the naïve to formative pluripotency transition. A, Venn diagram showing the overlap between FoxO1 targets (purple) and genes differentially expressed during WT differentiation (DEGs, log2FC N24 vs 2i ≤ −0.5 | ≥ 0.5, *p-adj.* ≤ 0.05, grey). The resulting *p-value* from the hypergeometric test of the overlap is shown. B, Volcano plot showing RNA-seq data from WT differentiation (N24 vs 2i). DEGs (as defined above) that are bound by FoxO1 are colour coded depending on whether they are bound only in 2i (green), only at N24 (light blue) or in both conditions (purple). Selected naïve and formative genes are indicated in the plot. For each quadrant, percentages (%) of FoxO1 2i-only, N24-only and 2i&N24 targets are shown. C, Expression dynamics of 2i-specific (green), N24-specific (light blue) and 2i&N24 (purple) FoxO1 targets measured across a 2 hours-resolved WT differentiation time course by RNA-seq (Lackner et al. 2021). Mean and confidence interval (CI) of log2FC values (relative to 2i) are shown. D, Genome browser snapshots showing FoxO1 ChIP signal in WT cells over selected formative core marker genes in 2i (green) and at N24 (light blue). Called peaks are also indicated in the plots (N24-only peaks: light blue, shared peaks: purple). E, Box plot showing the expression of exit factors (as defined in the text) that are upregulated during WT differentiation (log2FC > 0) in WT cells at N24, divided into FoxO1 N24 targets (light blue) or non-targets (grey), as measured by RNA-seq. Data is shown as log2FC relative to 2i. The resulting *p-value* from two-tailed Wilcoxon rank sum test is indicated in the plot. F, Venn diagram showing the overlap between FoxO1 targets (purple) and genes differentially expressed in *Pten* KO at N24 (DEGs, log2FC *Pten* KO vs WT ≤ −0.5 | ≥ 0.5, *p-adj.* ≤ 0.05, grey). The resulting *p-value* from the hypergeometric test of the overlap is shown. G, Box plot showing the expression of core formative genes (as defined in the text) in *Pten* KO cells at N24, divided into FoxO1 N24 targets (light blue) or non-targets (grey), as measured by RNA-seq. Data is shown as log2FC relative to WT. The resulting *p-value* from two-tailed Wilcoxon rank sum test is indicated in the plot.

**Supplementary Fig. 5.**
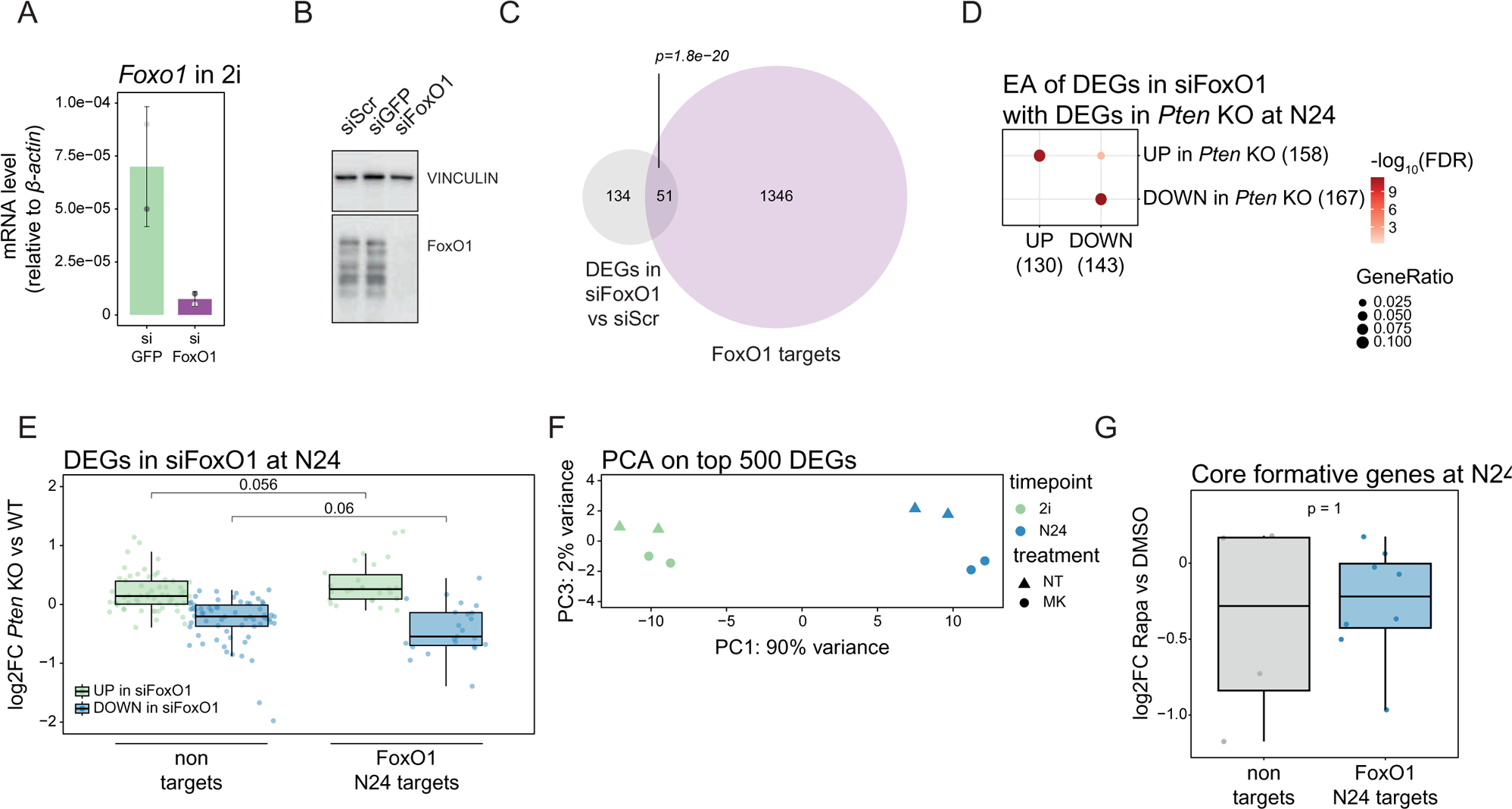
Interference with FoxO1 nuclear shuttling impairs the transition from the naïve to the formative GRN. A, *Foxo1* expression levels measured by RT-qPCR in WT cells transfected with control (siGFP, green) or siRNAs targeting FoxO1 (siFoxO1, purple). Mean and SD of n=2 independent experiments (depicted as distinct shades of grey) are shown. Expression was normalised to *β-actin*. B, Western blot analysis for FOXO1 expression in cells transfected with control (siGFP and siScr) or siRNAs targeting FoxO1 (siFoxO1). VINCULIN was used as a loading control. One representative of n=3 independent experiments is shown. C, Venn diagram showing the overlap between FoxO1 targets (purple) and genes differentially expressed in siFoxO1 at N24 (DEGs, *p-adj.* ≤ 0.2). The *p-value* from the hypergeometric test of the overlap is shown. D, Enrichment analysis (EA) of up-(UP) and down-regulated (DOWN) genes in siFoxO1 at N24 with upregulated (UP) or downregulated (DOWN) genes in *Pten* KO at N24. Dot colour and size indicate the FDR values (only FDR ≤ 0.05 are shown) and the GeneRatio (ratio between overlap size and the category size), respectively. E, Box plot showing the expression of up-(UP, green) or downregulated (DOWN, light blue) genes upon *Foxo1* knockdown (siFoxO1) in *Pten* KO cells at N24, divided into FoxO1 N24 targets or non-targets. Data is shown as log2FC relative to WT. The resulting *p-value* from two-tailed Wilcoxon rank sum test is indicated in the plot. F, PCA analysis based on the top 500 DEGs in RNA-Seq data of WT cells non-treated (NT, triangles) or after treatment with 1 μM MK-2206 (circles), in 2i (green) or at N24 (light blue). G, Box plot showing the expression of core formative genes in WT cells after Rapamycin treatment at N24, divided into FoxO1 N24 targets (light blue) or non-targets (grey). Data is shown as log2FC relative to DMSO-treated cells. The resulting *p-value* from two-tailed Wilcoxon rank sum test is indicated in the plot.

## Notes

### Competing Interest Statement

The authors have declared no competing interest.

